# Stage-Specific Transcriptome Landscape of Hepatocellular Carcinoma: Insights from Super and Poor Survivors with Prognostic Signature Identification

**DOI:** 10.1101/2024.11.26.625545

**Authors:** Xiao-Qian Xu, Hao Wang, Li-Chen Shi, Cheng Huang, Hong You, Ji-Dong Jia, You-Wen He, Yuan-Yuan Kong

**Affiliations:** National Clinical Research Center for Digestive Diseases, Beijing Friendship Hospital, Capital Medical University, Beijing, China; Clinical Epidemiology and EBM Unit, Beijing Clinical Research Institute. Beijing, China; Liver Research Center, Beijing Friendship Hospital, Capital Medical University, National Clinical Research Center for Digestive Diseases, Beijing, China; Department of Integrative Immunobiology, Duke University School of Medicine, Durham, NC 27710, USA

**Keywords:** Hepatocellular Carcinoma, Transcriptome, Stage-specific, Prognosis

## Abstract

**Background:** The prognosis for hepatocellular carcinoma (HCC) patients can vary significantly even within the same clinical stage. This study aims to characterize the stage-specific transcriptomic landscape of super survivors with HCC and identify a prognostic gene signature.

**Methods:** We analyzed data from The Cancer Genome Atlas (TCGA) for 76 matched super survivors (alive at 5 years) and poor survivors (deceased within 1 year) with HCC. Gene set enrichment analysis stratified by stage was conducted and a key prognostic gene signature was developed. The signature was then applied to the full TCGA cohort and independently validated in International Cancer Genome Consortium (ICGC) data.

**Findings:** Stage-specific transcriptomic profiling revealed distinct patterns in super survivors. Stage I and II showed positive enrichment in immune response pathways, while stage III exhibited enhanced catabolic activities and reduced glycolysis. Across all stages, cell cycle processes were less active in super survivors. A 19-gene signature, incorporating immune, metabolic, and cell cycle genes, accurately distinguished super from poor survivors with 91% accuracy (69/76). The signature reliably predicted overall survival in both the verification cohort (1-year, 3-year, and 5-year AUROCs of 0.82, 0.80, and 0.78) and the independent validation cohort (1-year and 3-year AUROCs of 0.80 and 0.83). Consistent AUROCs were observed across all tumor stages.

**Interpretation:** Our findings reveal a dynamic shift in HCC progression, with immune dominance in early stages and metabolic dominance in later stages, alongside reduced cell cycle activity across all stages. Integrating stage-specific transcriptomic profiles offers a promising approach to enhancing individualized management for HCC patients.

## Introduction

Hepatocellular carcinoma (HCC) prognosis is closely tied to tumor stage, with early-stage patients typically benefiting from curative treatments and achieving five-year survival rates exceeding 70%.^1^. In contrast, patients with advanced-stage HCC, treated primarily with systemic therapies, have a median survival of only 1 to 1.5 years^1^. However, even within the same clinical stage, patient outcomes can vary widely^2^. Approximately 7% to 10% of early-stage patients experience poor survival outcomes, while 3% to 10% of advanced-stage patients may survive beyond five years^3^. Previous studies associated these disparities with distinct tumor microenvironments and molecular activities linked to HCC progression, yet further investigation into the molecular factors that determine super and poor survival within each stage remains necessary.

Recent advances in transcriptome analysis have been instrumental in uncovering molecular subtypes, biomarkers, and gene expression patterns correlated with patient outcomes. Despite the development of numerous gene signatures for prognostic predictions, practical clinical utility remains elusive ^4^. Various deregulated pathways including those related to immune response, metabolism, hypoxia, cuproptosis, and autophagy, have been implicated in HCC prognosis ^5–8^. However, most existing gene signature are for the general HCC population without considering tumor stage, or they are specific to early or advanced stages. A comprehensive, stage-specific transcriptomic landscape comparing super and poor survivors is still underexplored, and uncovering such differences could offer significant prognostic insights for patients within the same stage.

Current molecular subclasses and gene signatures for prognosis are often developed using unsupervised clustering methods ^9,10^ or by comparing tumor tissues with adjacent non-tumor or normal tissues^11,12^, without fully leveraging core prognostic information like overall survival. We propose a more focused approach by comparing two extreme patient groups—super survivors and poor survivors—aiming to identify key gene signatures with the most critical prognostic significance. By concentrating on survival extremes, we hope to uncover genes with the strongest impact on patient outcomes, providing more accurate prognostic markers.

In this study, we characterized the transcriptomic landscape of super and poor survivors among HCC patients and identified stage-specific molecular alterations. We then developed a gene signature applicable across the entire patient population, aiming to improve the accuracy of overall survival predictions and contribute to more personalized treatment strategies for HCC.

## Materials and methods

### Data access and definitions

The gene expression data generated by RNA-Seq of cancer tissues, clinical and survival data of HCC patients were downloaded from The Cancer Genome Atlas Liver Hepatocellular Carcinoma (TCGA-LIHC) data (**Supplementary methods**)^13^. Cases were staged according to the American Joint Committee on Cancer (AJCC). Only HCC patients with RNA-Seq, clinical, and survival data were included in the final analysis. Informed consent was not applicable in this study, as the TCGA data was accessed in compliance with data-sharing policies, ensuring patient privacy and confidentiality.

Super survivors with HCC were defined as being alive at 5 years after HCC diagnosis, regardless of subsequent clinical outcomes. Poor survivors with HCC were defined as death ≤ 1 year since HCC diagnosis. To minimize confounding factors, poor survivors and super survivors were matched on sex (exact match) and age at HCC diagnosis (nearest matching) at 1:1 ratio.

### Identification of differentially expressed genes (DEGs)

Analysis on the identification of differentially expressed genes (DEGs) were performed in matched HCC patients with survival. The R package “DESeq” was applied to calculate expression alterations based on genes expression counts between super and poor survivors. Adjusted *p*-value < 0.05 and | log2(fold change) | >1 was set as the cut-off criteria for the identification of DEGs. Heatmap was generated with R package “pheatmap” based on the normalized log 2 (x+1) expression level of transcripts per million reads (TPM), which was converted from count format, with consideration of corresponding gene exon length.

### Gene set enrichment analysis (GSEA)

GSEA was utilized to analyze what biological processes were enriched in the gene rank between the super and poor survivors, both in all matched HCC patients and by tumor stage. The collection of annotated gene sets of Gene Ontology (GO) biological process and Hallmark pathways was defined as the reference gene sets and downloaded from Molecular Signatures Database (MSigDB)^14,15^. The R package “clusterProfiler” was applied at GSEA analysis.

### Infiltrating immune cells inferring

To identify the profile of infiltrating immune cells in HCC patients with super and poor survival, single-sample gene set enrichment analysis (ssGSEA) algorithm was performed through the “gsva” R package, with the TISIDB as the reference gene sets.^16^ The TISIDB database is an integrated repository designed to explore the interaction between tumors and the immune system, initiated by The University of Hong Kong, China. The database integrates the correlation matrix of genes and 28 immune cells, with data compiled from multiple sources, including high-throughput experiments, multi-omics data of pan-cancer, and other publicly available datasets.^17,18^ In our study, to show the different levels of immune cells between super and poor survivors, boxplot was presented. Boxplot edges indicated the 25th and 75th percentiles, and whiskers indicated nonoutlier extremes.

### Clinical significance analysis of key gene signatures in super and poor survivors

To identify key genes that best characterize super and poor survivors, all DEGs identified in the top 15 biology process that related to immune response, metabolism, and cell cycle were analyzed using cox regression model and selected based on Akaike information criterion (AIC), with the consideration of multicollinearity. Time-dependent area under receiver operating characteristic curves (AUROC) of selected gene sets were calculated and plotted in matched super and poor survivors. Kaplan Meier curves was performed to estimate the survival difference between the low-risk and high-risk groups, classified by the cutoff at the maximal Youden index. *P* values for survival curves were determined by use of the log-rank test. In matched super and poor survivors, time-dependent AUROC in predicting 5-year overall survival were calculated through utilizing package “timeROC” in R.

### Validation of identified key gene sets in total HCC patients from TCGA and ICGC

The identified key gene sets from super and poor survivors, alone and combined with key clinical factors, were validated in total HCC patients from both TCGA-LIHC and International Cancer Genome Consortium (ICGC), the project of Japan – Liver Cancer (Hepatocellular carcinoma; virus associated (**Supplementary Methods**)^19^. Cases were staged according to the tumor-node-metastasis staging proposed by Liver Cancer Study Group of Japan.

Missing data on clinical factors were imputed through predictive mean matching for continuous variables, and through logistic regression or polytomous logistic regression for categorical variables. Time-dependent AUROC in predicting 1-year, 3-year, and 5-year overall survival were calculated. Cutoffs of the low-risk, intermediate-risk, and high-risk groups were determined by the highest and lowest 20% quantiles of the predicted probabilities in total HCC patients from TCGA and ICGC.

### Statistical analysis

Statistical analyses were performed using R, version 3.5.0 (R Foundation for Statistical Computing). Two-tailed P < 0.05 was considered statistically significant, if not specified specifically. Descriptive analysis was shown in **Supplementary Methods**.

## Results

To investigate the transcriptional landscape of HCC tumors in super and poor survivors, we identified 363 HCC patients with survival and RNAseq data in TCGA-LIHC database. A total of 59 and 40 HCC patients met super and poor survival criteria, respectively. Based on the matching criteria of sex and age, 38 super and 38 poor survivors were matched and analyzed for the identification of DEGs and key gene signatures in a discovery phase **(Figure 1)**. The identified gene signatures for disease prognosis were then verified in the total 363 HCC patients and further validated in an independent dataset consisting of another 243 HCC patients with survival and RNAseq data from ICGC.

**Figure 1.**
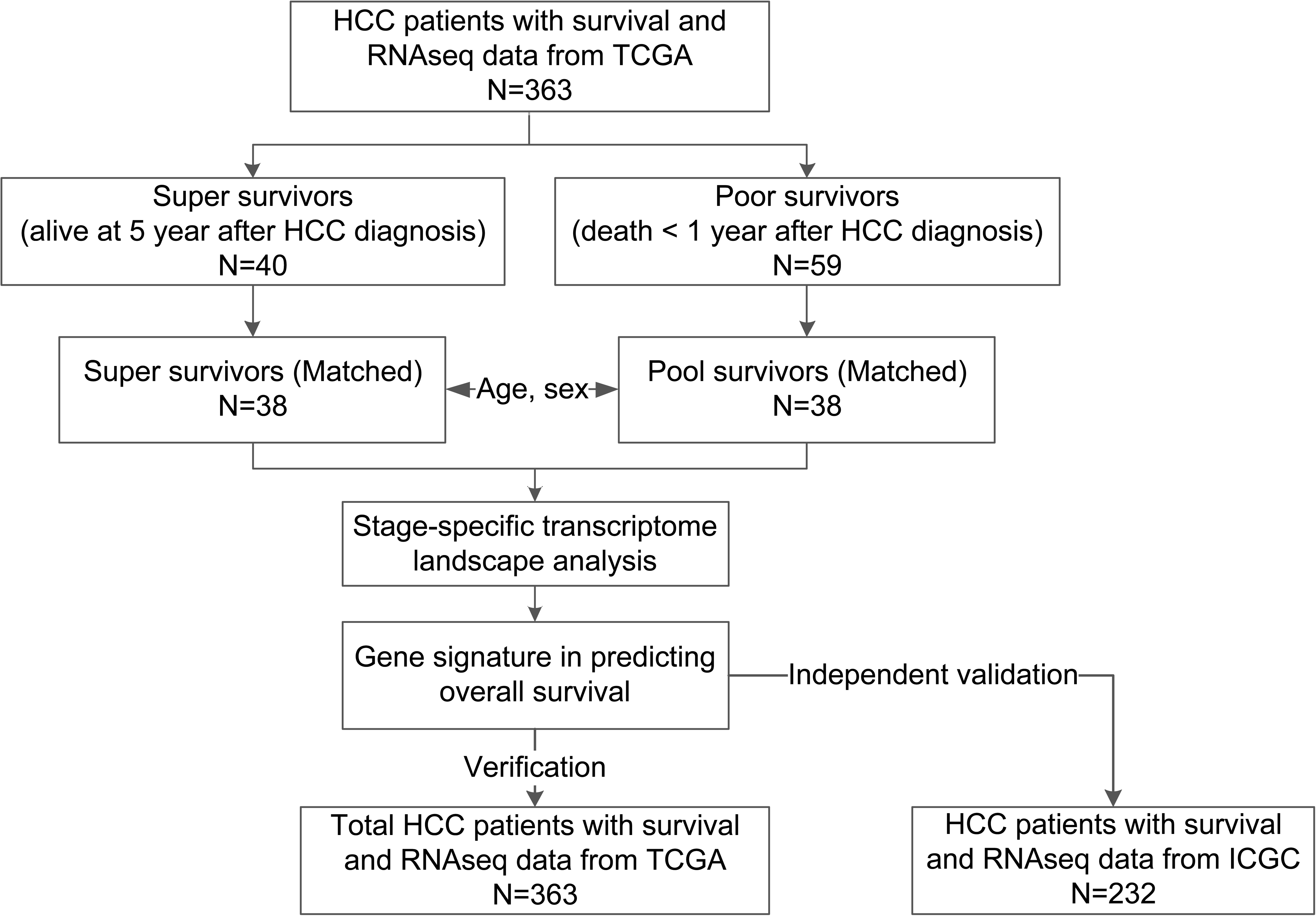
Flowchart.

### Clinical profiles of the HCC patients with super and poor survival

The demographic, pathologic, and biochemical characteristics in the HCC patients from TCGA and matched super and poor survivors were described in **Supplementary Table 1**. The median age of total HCC patients were 61 years old, with 67.49% being males. A total of 85 patients (25.1%) had stage [ and above. As expected, in the matched super and poor survivors, more severe tumor stage (*p*=0.034) and pathologic T stage (*p*=0.020) were identified in poor survivors. The median overall survival (mOS) in the total HCC patients was 4.64 (95%CI: 3.82, 6.93) years and the median follow-up time was 2.27 (95%CI:1.98, 2.71) years. The mOS of the poor survivors was 0.32 (95%CI: 0.18, 0.76) years. The mOS of the super survivors could not be calculated since the survival rate was higher than 50% until the end of follow-up, with the median follow-up time at 6.6 (95%CI: 6.3, 7.5) years.

### DEG identification and enrichment analysis in HCC super survivors

In the matched super and poor survivors, a total of 684 DEGs were identified, with 252 DEGs showing higher expressions in super survivors and 432 DEGs showing lower expressions in super survivors (**Figure 2A**). Detailed information on the fold change and *p* values of the top 200 DEGs were presented in **Supplementary Table 2**. A distinct distribution pattern of high-expression on the 252 DEGs and low-expression on the 432 DEGs were presented in the HCC super versus poor survivors (**Figure 2B)**. These data identified distinct mRNA gene expression patterns between super and poor survivors.

**Figure 2.**
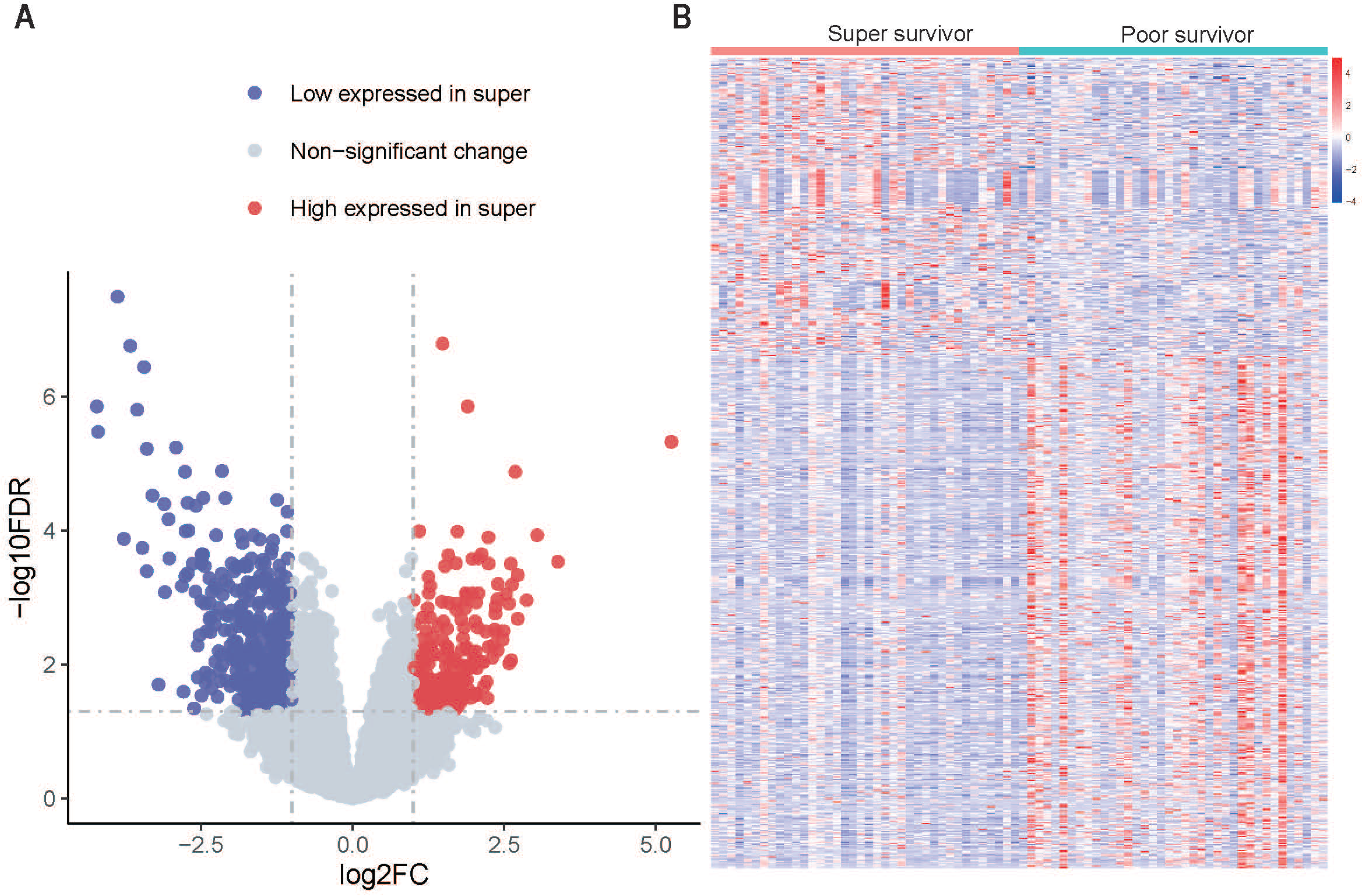
Transcriptomes in matched super and poor survivors with HCC. (A) Volcano plot of differently expressed genes in super versus poor survivors. (B) Heatmap of differently expressed genes in super versus poor survivors. Columns display the samples of super and poor survivors and rows indicate the up-regulated and down-regulated DEGs.

### GSEA enrichment analysis in HCC super survivors

To analyze the biological processes and pathways associated with HCC patient survival, we performed GSEA analysis based on the rank of the fold change on gene expressions in super versus poor survivors with GO terms and Hallmark pathways.

The top 15 GO terms and hallmark pathways were shown in **Figure 3A**. GO biology process GSEA identified several immune related processes, involving both B and T lymphocyte-mediated adaptive immunity, that was significantly up-regulated in super survivors. Different from GO process, Hallmark pathways have shown enrichment in metabolism pathways, including upregulated oxidative phosphorylation, bile acid, fatty acid and xenobiotic metabolism and adipogenesis as well as down-regulated glycolysis. Cell cycle related processes or pathways were consistently down-regulated in super survivors as found through GO process and Hallmark pathway (**Figure 3A**).

**Figure 3.**
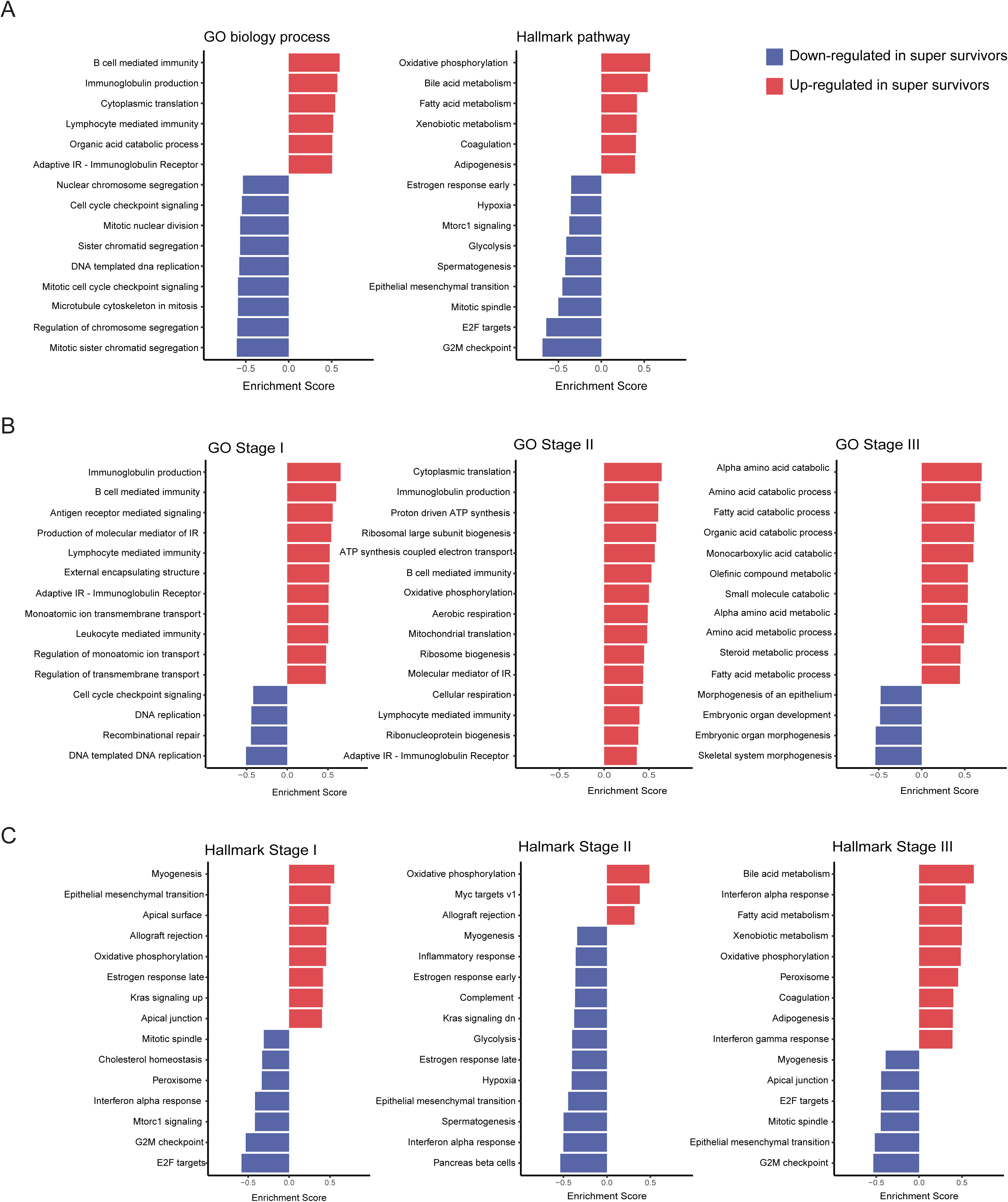
The biological processes and pathways enriched in matched super and poor survivors with HCC. The top 15 significantly GSEA enrichment of GO terms in biological processes and Hallmark pathways, ranked by *p*-value and normalized enrichment score in all matched super and poor survivors (A), by stage using GO (B), and by stage using Hallmark (C). GO, gene ontology. GSEA, gene set enrichment analysis. IR, immune response

GSEA enrichment analysis by tumor stages further demonstrated that immune related GO biological process was mainly enriched in HCC tumors from super survivors at tumor stage [ and [, whereas the metabolism pathways was mainly enriched in super survivors at tumor stage [ (**Figure 3B**), with the later also supported in Hallmark GSEA results (**Figure 3C**). More specifically, amino acid, fatty acid, organic acid and monocarboxylic acid catabolic process were up-regulated in tumors of the super survivors at stage [. In super survivors at tumor stage [, both immune related and metabolic processes, including oxidative phosphorylation and ATP synthesis were up-regulated (**Figure 3B and 3C**).

### Identification of key gene signature involved in immune responses in HCC super survivors

Since the immunological processes were significantly up-regulated in HCC tumors from the super survivors, the levels of infiltrating immune cells from RNAseq data were inferred using available resource^16^. The analysis revealed 19 distinct immune cell subsets in HCC tumors (**Figure 4A**). Consistent with GSEA enrichment showing both B and T lymphocyte involvement in super survivors, activated B cells and activated CD8^+^ T cells were significantly higher in HCC tumors in super survivors at all stages compared to those in poor survivors (**Figure 4A and 4B**). Immature B cell subset in super survivors diagnosed at tumor stage [ was significantly higher than that in poor survivors (**Figure 4B**). Monocyte subset was significantly higher only in super survivors diagnosed at tumor stage [ while three immune cell subsets, eosinophils, T helper 1 (Th1) and T helper 2 (Th2) cells, were significantly higher and regulatory T cells (Tregs) were lower in HCC tumors from super survivors diagnosed at tumor stage [ (**Figure 4B**).

**Figure 4.**
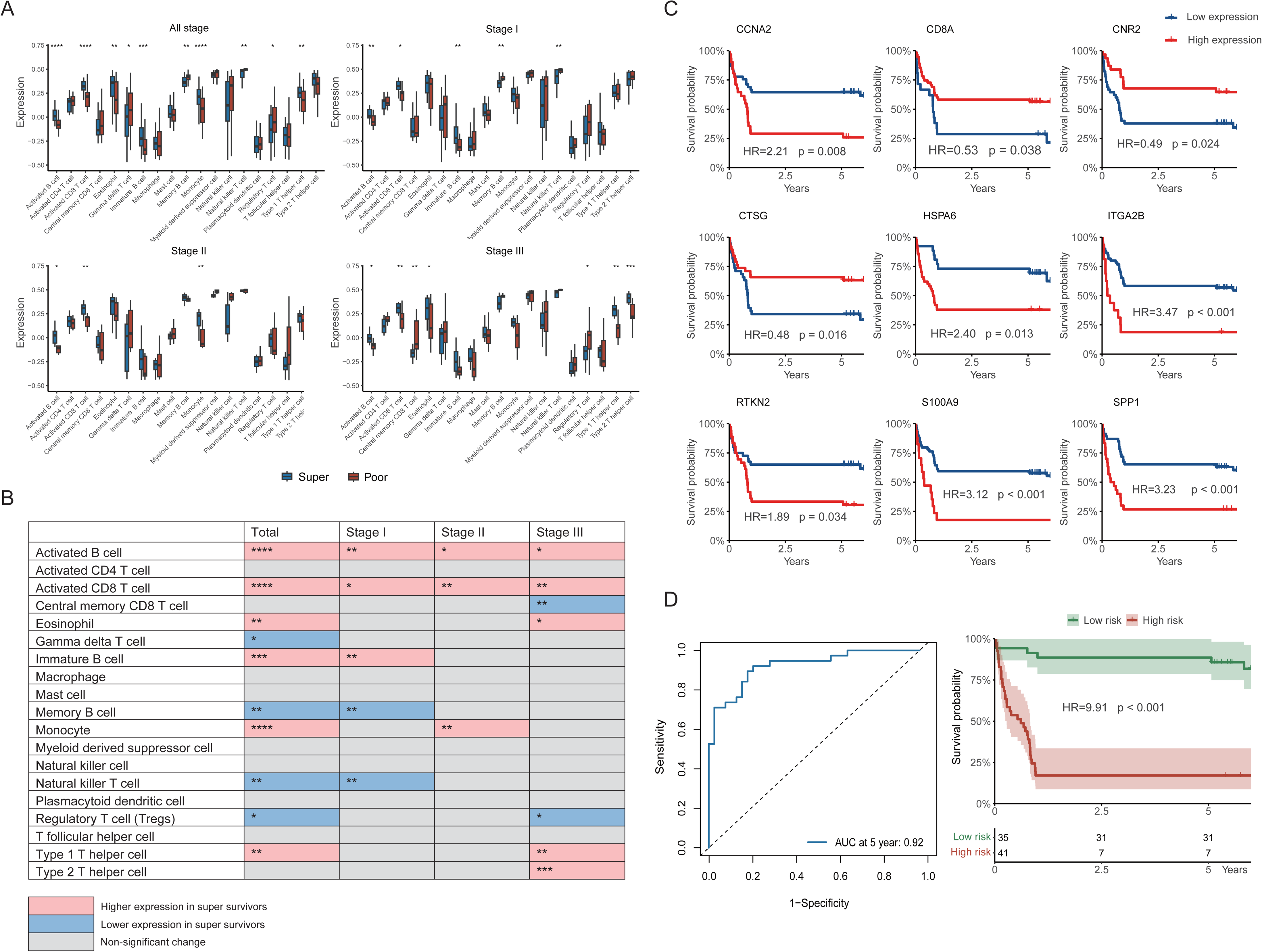
Key gene signature involved in immune response across super and poor survivors with HCC. (A) Bar plots of ssGSEA-inferred expressions of different immune cell types across super and poor survivors. (B) Characterization of inferred immune cell types in all matched super and poor survivors at different stages. (C) Kaplan-Meier survival curves of the individual key DEGs for inferring immune cell types. Cutoffs for low-risk and high-risk groups were determined by maximal youden index. (D) AUROC and Kaplan-Meier survival curves of gene sets involved in immune response.

To identify a key gene signature in the HCC super survivors, 35 DEGs related to immune pathways identified from matched super and poor survivors were used as the foundation for further analysis. After stepwise exclusion by multicollinearity and selection with AIC in Cox regression model, 9 genes involving in immunological processes were identified as potential components of a key gene signature panel, including CCNA2, CD8A, CNR2, CTSG, HSPA6, ITAG2B, RTKN2, S100A9 and SPP1 (**Supplemental Table 2**). All 9 genes individually showed significant HR varying from 3.47 to 0.48(**Figure 4C**). Combining the 9 immune related genes into a single signature panel showed a 5-year AUROC at 0.92 for the 76 super and poor survivors (**Figure 4D**). The immune gene signature panel stratified the 76 HCC patients at HR of 9.91 (p<0.001) between the two groups (**Figure 4D**). As shown in **Supplementary Figure 1**, 11 patients were misclassified using this immune gene signature, including 7 super survivors stratified into the high-risk group and 4 poor survivors stratified into the low-risk group. Taken together, the above analysis demonstrates that the immune gene signature alone can accurately predict the survival probability of 86% (65/76) of the HCC patients and suggests a need to incorporate other related gene signatures.

### Identification of key gene signature involved in metabolism in HCC super survivors

A total of 17 metabolism related processes or pathways encompassing 702 genes were identified from GSEA analysis in total matched super and poor survivors as well as in super survivors diagnosed at tumor stage [ (**Figure 3A and B**). Finally, 15 DEGs were expressed significantly lower and 10 DEGs significantly higher in super survivors comparing to poor survivors (**Figure 5A**).

**Figure 5.**
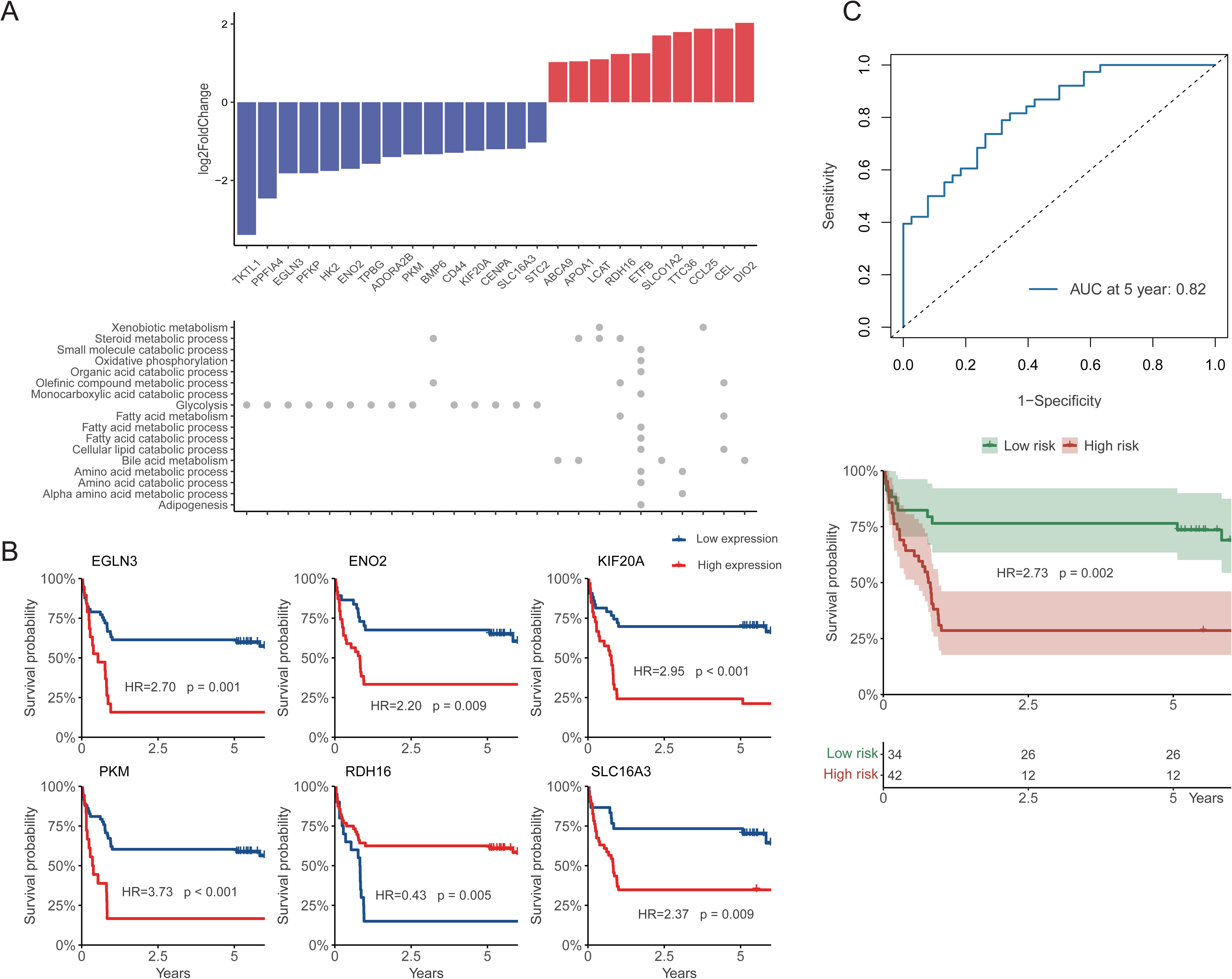
Key gene signature involved in metabolism across super survivors and poor survivors with HCC. (A) Characterization of all DEGs involved in metabolism across super and poor survivors. (B) Kaplan-Meier survival curves of the individual key DEGs involved in metabolism. Cutoffs for low-risk and high-risk groups were determined by maximal youden index. (C) AUROC and Kaplan-Meier survival curves of gene sets involved in metabolism.

After stepwise exclusion by multicollinearity and selection with AIC in Cox regression model, 6 genes related to metabolic process were identified, including EGLN3, ENO2, KIF20A, PKM, RDH16, and SLC16A3 (**Supplemental Table 2**). Except RDH16 whose high expression was associated with a lower risk (HR=0.43), the other 5 genes individually showed significant higher risk with HR varying from 2.20 to 3.73 in Kaplan-Meier curves plot (**Figure 5B**).

Combining the 6 metabolic related genes into a single signature panel showed a 5-year AUROC at 0.82 (**Figure 5C**). When used to stratify the 76 HCC patients, the metabolism signature yielded a HR of 2.73 (p=0.002) for survival probability between the high-risk and low-risk groups (**Figure 5C**). A total of 19 HCC patients were misclassified, including 12 super survivors stratified into the high-risk group and 7 poor survivors stratified into the low-risk group (**Supplementary Figure 1**), resulting a 75% (57/76) accuracy. This analysis suggests that immune gene signature is superior to the metabolism gene signature when used alone.

### Identification of key gene signature involved in cell cycle in HCC super survivors

Our analysis also identified a total of 14 cell cycle related processes or pathways that encompassed 553 genes from GSEA when comparing super versus poor survivors **(Figure 3**). Among the 553 genes, 49 were expressed at lower levels in super survivors comparing to those in poor survivors (**Figure 6A**). After stepwise exclusion by multicollinearity and selection with AIC in Cox regression model, 5 genes related to cell cycle process were identified, including CD44, CXCL1, SPOCK1, SPP1, and TTK (**Supplemental Table 2**). All 5 gene individually showed significant HR varying from 2.66 to 4.56 in Kaplan-Meier curves plot (**Figure 6B**). Remarkably, almost all patients with high expressions of CD44 and TTK died within 1 year (**Figure 6B**). Combining the 5 cell cycle-related genes into a single gene signature showed a 5-year AUROC at 0.82 (**Figure 6C**).

**Figure 6.**
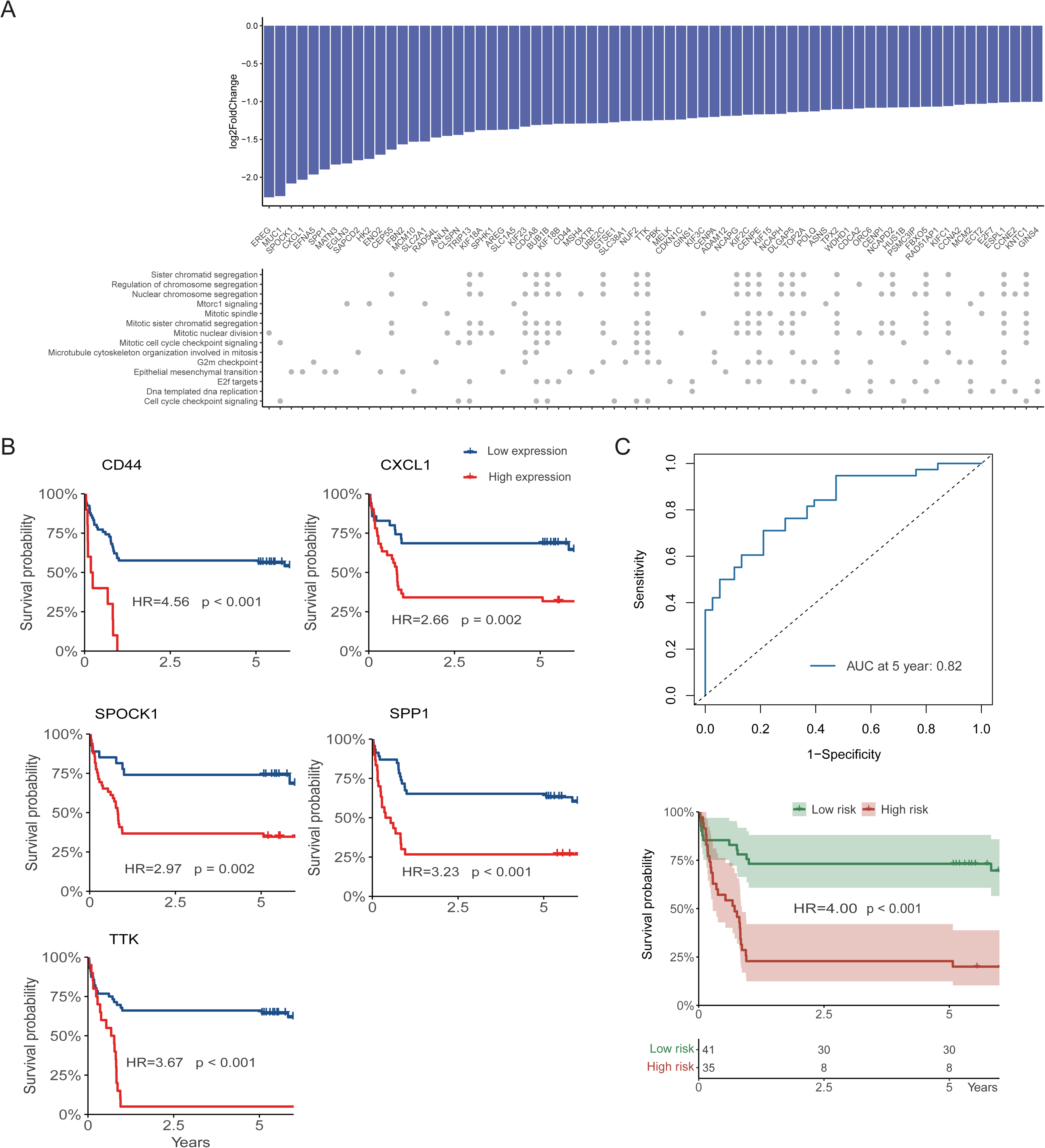
Key gene signature involved in cell cycle across super and poor survivors with HCC. (A) Characterization of all DEGs involved in cell cycle across super and poor survivors. (B) Kaplan-Meier survival curves of the individual key DEGs involved in cell cycle biology process. Cutoffs for low-risk and high-risk groups were determined by maximal youden index. (C) AUROC and Kaplan-Meier survival curves of gene sets involved in cell cycle.

When used to stratify the 76 HCC patients, the cell cycle signature yielded a HR of 4.00 (p<0.001) for survival probability between the high-risk and low-risk groups **(Figure 5C).** A total of 19 HCC patients were misclassified, including 8 super survivors stratified into the high-risk group and 11 poor survivors stratified into the low-risk group (**Supplementary Figure 1**), resulting a 75% (57/76) accuracy. This analysis suggests that immune gene signature is superior to the cell cycle gene signature when used alone.

### Prognostic performance of a combined gene signature for HCC patient survival

While the immune gene signature showed good performance in discriminating super and poor survivors, metabolism and cell cycle gene signatures might help to rectify some of the misclassified super or poor survivors (**Supplementary Figure 1**). We then constructed a combined gene signature panel involving all three biological processes, consisting of 19 genes due to that SPP1 was identified in both immune and cell cycle related gene signatures.

The 19 gene signature panel showed a 5-year time-dependent AUROC at 0.96 in the 76 HCC patients (**Figure 7A**). When stratifying the survival probabilities into low-risk and high-risk, the combined gene signature reduced the misclassified HCC patients to 7, including 4 super and 3 poor survivors, resulting in a 91% (69/76) accuracy (**Supplementary Figure 1**). The HR for survival probability between the high-risk and low-risk groups was improved to 16.24 (p<0.001).

**Figure 7.**
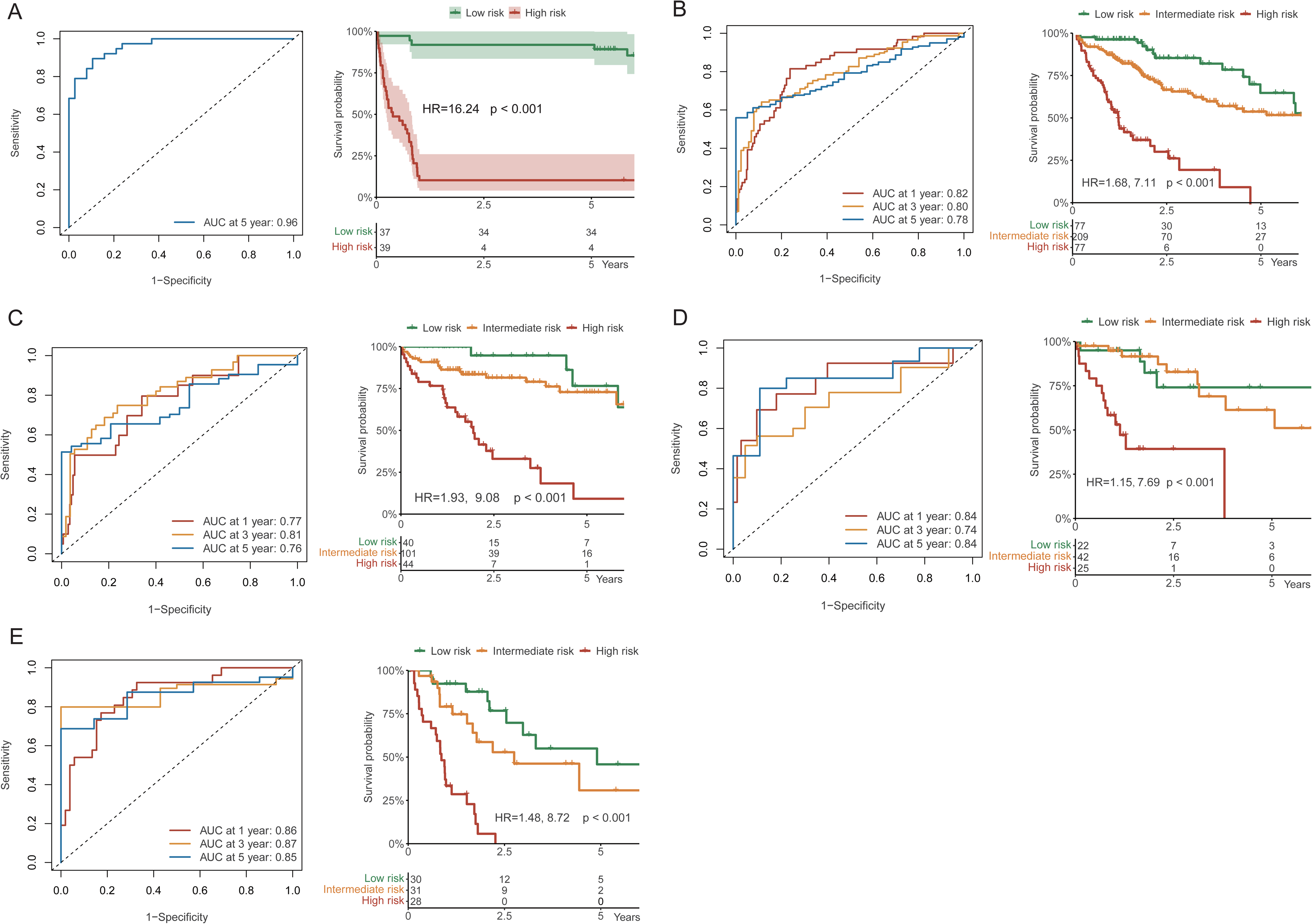
Clinical performance of the combined gene signature. AUROC and Kaplan-Meier survival curves of the combined gene signature in predicting the overall survival in matched super and poor survivors with HCC (A), total HCC patients from TCGA (B), and HCC patients from TCGA in stage [ (C), stage [ (D) and stage [ (E).

To verify the clinical performance of the combined gene signature, its predicating capability was evaluated in the total 363 HCC patients from TCGA-LIHC. The 1-year, 3-year, and 5-year time-dependent AUROCs were 0.82, 0.80 and 0.78, respectively (**Figure 7B).** Patients stratified into high-risk and intermediate-risk groups showed 7.11-fold and 1.68-fold higher risks on total death than patients stratified into low-risk groups (**Figure 7B)**. The performance verified by tumor stages revealed relatively higher AUROCs in stage □ and III than AUROCs in stage □ (**Figure 7C, 7D, and 7E**). The HR for survival probability in the high-risk and intermediate-risk groups compared with low-risk groups was 9.08 and 1.93 at stage □ (p<0.001), 7.69 and 1.15 at stage □ (p<0.001), and 8.72 and 1.48 at stage □ (p<0.001).

To further enhance the predicting capability of the combined gene signature in HCC patient survival, we tested the addition of 5 clinical factors. Among 11 clinical factors, 5 including age at HCC diagnosis, BMI, tumor stage, pathologic T stage, and AFP levels were included based on stepwise AIC criteria. Addition of these clinical factors yielded 1-year, 3-year, and 5-year time-dependent AUROCs of 0.85, 0.84 and 0.87 respectively for the total patient population (**Supplementary Figure 2A**). This improvement was most pronounced in patients at stage □, with the 1-year, 3-year, and 5-year time-dependent AUROCs increased from 0.77, 0.81, and 0.76 to 0.86 (increased by 11.69%), 0.89 (9.9%), and 0.85 (11.84%) (**Supplementary Figure 2B**). The Kaplan-Mier curves stratified by gene signature and clinical factors were similar to that stratified by gene signature alone. However, the HRs in the high-risk and intermediate groups were improved to 42.33 and 6.58 in patients at stage [ (**Supplementary Figure 2D**). Taken together, these data demonstrate that the combined gene signature provides an acceptable prognosis capability, and addition of 5 clinical factors further enhances the gene signature’s power.

### Independent validation of the combined gene signature in HCC patients

To validate the above gene signature in an independent HCC patient population, we obtained data from ICGC database on 232 HCC patients from Japan. The median age at HCC diagnosis was 68.5 (95%CI: 62, 74) years old. Males accounted for 73.7% of the patients, which were similar to data from TCGA. More patients were diagnosed at stage [ and stage [ in ICGC (45.7% and 38.8% respectively) compared to data from TCGA (24.8% and 25.1 % respectively). The median follow-up time was 2.46 (2.30, 2.63) years. The 1-year and 3-year survival probabilities were 92.6% (95%CI: 89.2%, 96.0%) and 81.5% (95%CI: 76.1%, 87.3%).

The combined gene signatures showed time-dependent AUROC at 1-year and 3-year at 0.80 and 0.83 (**Figure 8A**). Patients stratified into high-risk and intermediate-risk groups showed a 32.40-fold and 6.83-fold higher risk on total death compared with patients stratified into low-risk groups (p<0.001) (**Figure 8A).** The patients were also stratified according to their tumor stages. For patients diagnosed at stage [, high AUROCs at 1-year (AUROC=0.99) and 3-year (AUROC=0.86) follow-up were achieved (**Figure 8B)**. HRs for high-risk and intermediate-risk groups could not be calculated due to that no patients stratified into low-risk group died until the end of follow-up. For patients diagnosed at stage [, AUROCs were at 0.74 at 1-year and 0.81 at 3-year follow-up (**Figure 8C)**.

**Figure 8.**
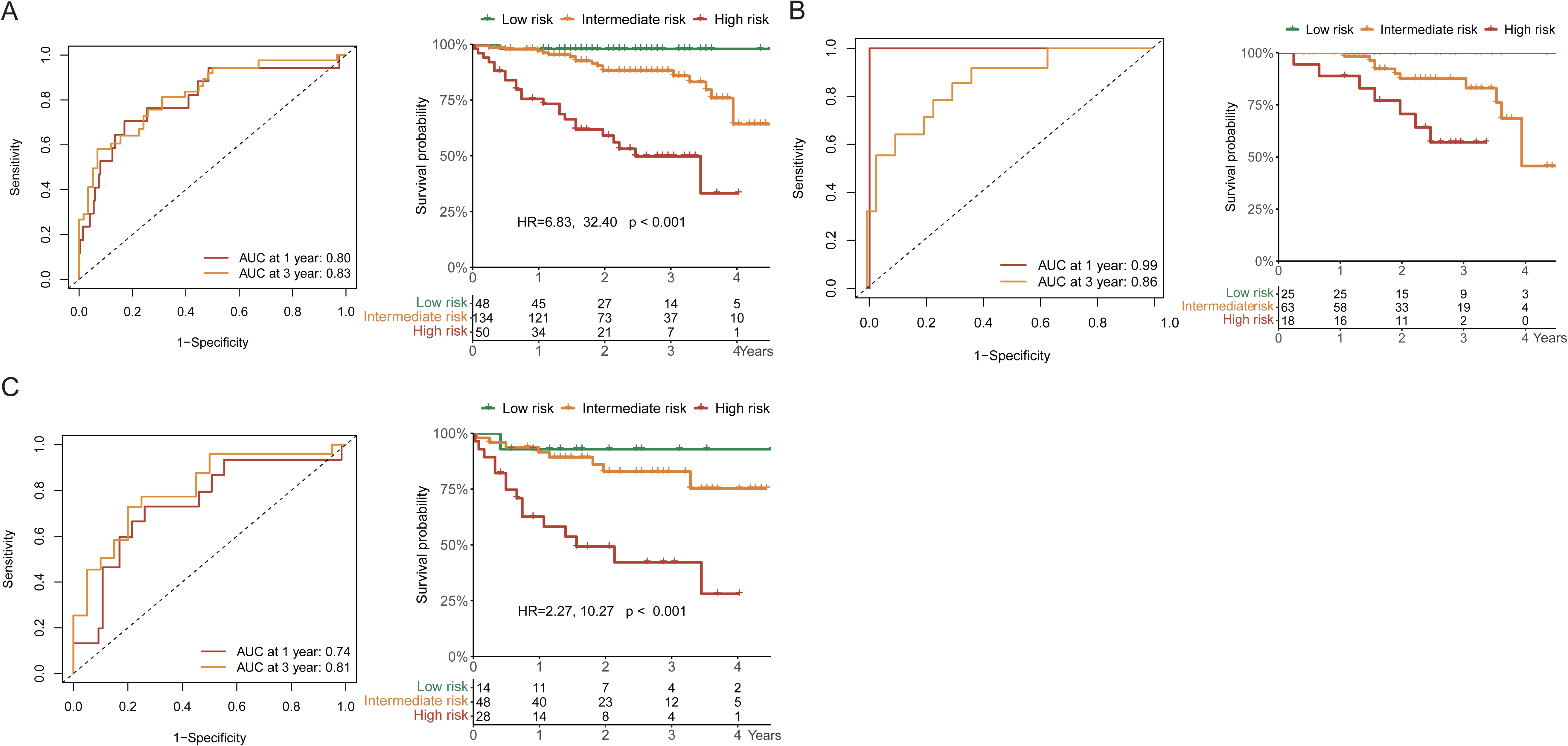
Clinical performance of combined gene signatures in ICGC. AUROC and Kaplan-Meier survival curves of combined gene signatures in predicting the overall survival in total HCC patients from ICGC (A), and HCC patients from ICGC in stage [ (B) and stage [ (C). Stage [ were not plotted as there’s only one death in this group.

Taken together, the above analysis suggests that the combined gene signature without the addition of clinical factors accurately predict the survival probability when applied to an independent validation cohort. We also extracted 14 prognostic gene signatures for HCC overall survival from the literature, each with AUROCs reported and external validation (**Supplementary Table 4)**. These AUROCs ranged from 0.65 to 0.80 across various time points (1-, 2-, 3-, 4-, and 5-year survival) in both discovery and validation cohorts. Notably, none of these signatures report stage-specific performance. In contrast, our 19-gene signature consistently achieved AUROC values around 0.80 across different time points and tumor stages, indicating enhanced predictive power for HCC overall survival.

## Discussions

Our study revealed a stage-specific transcriptome landscape in HCC super survivors, marked by early-stage immune responses, later-stage metabolic shifts, and consistently reduced cell cycle activities across all stages. Furthermore, we identified a 19-gene signature that accurately distinguished super survivors from poor survivors, reflecting immune, metabolism, and cell cycle pathways. This gene signature reliably predicted overall survival across both verification and independent validation cohorts, demonstrating prognostic accuracy across different tumor stages.

The enrichment of immune responses in stages I and II, along with the increasing dominance of metabolism-related pathways in later stages, supports our hypothesis that the biological pathways contributing to survival vary by tumor stage. This temporal shift, where immune activities predominate early but decline as metabolic processes take over in later stages, underscores the heterogeneity of the tumor microenvironment over time. Tumor-driven immunoediting—transitioning from immune surveillance to immune tolerance—may explain this shift ^20^. Our ssGSEA results further corroborate this by identifying stage III-specific prognostic relevance of regulatory T cells (Tregs), Th1, and Th2 cells. Despite the reduced immune activity in advanced stages, the sustained importance of activated B cells and CD8+ T cells across all stages suggests immune processes continue to play a role, even as metabolism-related pathways become increasingly prominent. At later stages, reduced glycolysis and increased catabolic activity, as predicted by our signature, point to better survival outcomes. This finding aligns with the "Warburg effect", where excessive glycolysis leads to acid accumulation, contributing to hepatocyte damage and cell death^21,22^. Further research into these mechanisms could open pathways for novel therapies targeting metabolic dysfunction in advanced HCC. ^21^ Throughout all stages, the persistent activity of cell cycle pathways highlights their continuous role in HCC progression. This dynamic shift—from immune dominance in early stages to metabolic dominance in later stages, with cell cycle involvement throughout—emphasizes the evolving nature of HCC immune pathophysiology.

The integration of immune response, metabolism, and cell cycle pathways in a prognostic gene signature realized higher AUROC and lower misclassification rate than any individual pathway alone in super and poor survivors. This suggests that each pathway contributes uniquely to tumor behavior in the dynamics of HCC progression and signature from all three components yields complementary prognostic information. When the combined gene signature was applied to total patients, a relatively high prognostic performance was achieved, with 1-, 3-, 5-year time dependent AUROC at 0.82, 0.80 and 0.78. The addition of clinical factors has further improved the 1-year, 3-year, and 5-year time-dependent AUROC to 0.85, 0.84 and 0.87 in total patients. Many of the signatures developed for HCC prognosis have highlighted deregulation of specific molecular pathways. For instance, an mRNA signature related to immunity reported 3-year and 5-year AUROCs at 0.72^6^, while a DNA damage response-related prognostic model showed a decline in performance over time, with the AUROC decreasing from 0.80 at 1 year to 0.69 at 3 years^23^. Similarly, an immune- and metabolism-related gene signature achieved a 3-year AUROC of 0.79, but its performance dropped to 0.71 and 0.69 at 3 and 5 years, respectively^11^. In contrast to these studies, which focus on specific pathways, our gene signature offers a broader and dynamic biological perspective. The consistent AUROC values over time highlight the reliability of our signature for both short- and long-term prognostic predictions.

When the signature was verified by tumor stage, the prognostic performance was relatively lower in patients at stage [ than patients at stage [ and [ when using the gene signature alone. This observation aligns with the current diagnostic difficulty mainly in early-stage HCC lesions.^24^ A possible explanation is that in early-stage HCC, molecular characteristics may still closely resemble normal tissue architecture. Nevertheless, when the clinical factors were added, an apparent improvement on AUROC at around 10% in stage [ was observed, achieving levels comparable to those seen in stage [ and [. More importantly, although AUROC represent a more comprehensive indicator, the capacity for risk stratification is more informative for clinical decision-making^25^. The developed gene signature, whether applied independently or in combination with clinical factors, effectively identified high risk patients at every tumor stage. This reliable risk stratification highlights its potential clinical utility in tailoring patient management strategies. The increasing affordability and accessibility of transcriptome sequencing technologies also enhance the feasibility of incorporating gene signatures into routine clinical practice.

The validity of our gene signature was further reinforced by the independent validation in virus-associated HCC patients from Japan, where it demonstrated strong prognostic performance, even in the absence of clinical factors. Compared to non-viral HCC, HBV and HCV infections are usually associated with early molecular alterations involved in the induction of chronic inflammation during viral oncogenesis (e.g., HBV DNA integration, chronic HCV infection-induced oxidative stress).^24^ As we identified from matched super and poor survivors, the immune related gene signature showed more predominant predictability than other two pathways in this model. Consequently, this gene signature, which is closely linked to immune responses, is likely to be even more predictive in virus-associated HCC, despite being developed from a cohort with diverse etiologies.

One limitation of the study is the absence of treatment regimens received by patients during follow-up. However, the observation period of all HCC patients analyzed occurred before 2020, predating the approval of the combination of atezolizumab and bevacizumab as a first-line treatment for advanced HCC ^26^. The therapies were likely similar across patients at comparable stages based on standard treatment guidelines. Another limitation of this study is the relatively high censoring rate in the TCGA data. By first comparing super vs. poor survivors, the study ensures that the identified genes are highly relevant for predicting outcomes and less susceptible for censor during follow-up. Additionally, the lack of etiology in the discovery cohort limits the ability to draw specific insights into the causes of HCC. However, the study’s objective was to develop a gene signature that applies broadly, irrespective of etiology. The signature’s performance was further validated in an independent cohort of virus-associated HCC patients, reinforcing the robustness and generalizability of the findings.

In conclusion, our study highlights the dynamic, stage-specific transcriptome landscape of HCC and identifies a robust 19-gene signature that integrates immune, metabolic, and cell cycle pathways. This signature offers reliable prognostic predictions across patient groups and tumor stages, supporting its potential clinical utility in personalizing HCC management. Future research into the mechanistic underpinnings of these pathways could provide valuable insights for developing targeted therapies in HCC.

## Acknowledgments

The authors acknowledge the participants and their families who contributed to make this study possible.

## Abbreviation

AIC: Akaike information criterion;
AUROC: area under receiver operating characteristic curves;
DEG: differentially expressed gene;
GO: Gene Ontology;
GSEA: gene set enrichment analysis;
HCC: hepatocellular carcinoma;
ICGC: International Cancer Genome Consortium;
ssGSEA: single-sample gene set enrichment analysis;
TCGA: The Cancer Genome Atlas;
TCGA-LIHC: The Cancer Genome Atlas Liver Hepatocellular Carcinoma

## Data availability statement

Publicly available datasets were analyzed in this study. This data can be found from The Cancer Genome Atlas (TCGA) (https://portal.gdc.cancer.gov/) and International Cancer Genome Consortium (ICGC) (https://icgc.org/) databases.

