## Supplementary material for "Stage-Specific Transcriptome Landscape of Hepatocellular Carcinoma: Insights from Super and Poor Survivors with Prognostic Signature Identification"

**Supplementary Materials**

**Supplementary Methods.**

TCGA-LIHC is a part of the TCGA project, which aims to collect and analyze large-scale cancer genomics. It is jointly supported and managed by the National Cancer Institute and the National Human Genome Research Institute of the U.S. National Institutes of Health^[1]^. Specifically, TCGA-LIHC focuses on HCC, regardless of etiology. Data in the TCGA-LIHC dataset was deposited by a collaborative effort involving research institutions and hospitals mainly in United States. The samples represent a diverse group of patients in terms of demographics, tumor types, and stages. It follows strict protocols for data collection, processing, and quality control. The TCGA-LIHC data has been cited in numerous high-impact studies, making it one of the most trusted HCC databases in biomedical research^[2-5]^.

The ICGC Japan-Liver Cancer database is part of the ICGC program, which was launched to coordinate large-scale cancer genomics from 50 different cancer types and/or subtypes across the globe, with data utilized in numerous literature ^[1,6,7]^. The Japan-Liver Cancer project, specifically, focuses on virus associated HCC in Japanese patients. It included the RNA-Seq expression and corresponding clinical information of 243 liver cancer samples and 202 cancer adjacent tissues from 232 patients^[8,9]^. In the study, only the first sample from each patient was used for validation.

Descriptive analysis for demographic factors and biochemical measurements available in TCGA and ICGC were performed. Continuous variables were presented as means and standard deviations (SD) or medians and interquartile ranges according the data distribution. For comparisons between groups, the student's t-test and the Mann-Whitney U-test were applied for normal and non-normal data, respectively. Categorical variables were presented as frequencies and percentages and compared using the Chi-square test or Fisher’s exact test.

**Supplementary Table 1. Clinical features of age and sex matched super and poor survivors with HCC**

|  | Total patients  (N=363) | Matched poor  (N=38) | Matched super  (N=38) | P value |
| --- | --- | --- | --- | --- |
| Age, years | 61(51, 69) | 67(61, 72) | 59(53, 68) | 0.0863 |
| Male (%) | 245 (67.5) | 26 (68.4) | 26 (68.4) | >0.9999 |
| BMI | 24.5 (21.8, 28.7) | 22.4 (20.4, 25.2) | 25.5 (23.3, 29.7) | 0.0197 |
| Race (%) |  |  |  | 0.2269 |
| American Indian/Alaska Native | 1 (0.3) | 0 | 0 |  |
| Asian | 155 (43.9) | 19 (51.4) | 18 (48.7) |  |
| Black/African American | 17 (4.9) | 3 (8.1) | 0 |  |
| White | 180 (51.0) | 15 (40.5) | 19 (51.4) |  |
| Family history of cancer (%) | 110 (35.0) | 9 (25.7) | 13 (39.4) | 0.3016 |
| Tumor grade (%) |  |  |  | 0.6783 |
| G1 | 55 (15.4) | 7 (18.4) | 7 (18.9) |  |
| G2 | 175 (48.9) | 17 (44.7) | 20 (54.1) |  |
| G3+ | 128 (35.8) | 14 (36.8) | 10 (27.0) |  |
| Tumor stage (%) |  |  |  | 0.0342 |
| Stage I | 170 (50.2) | 13 (35.1) | 23 (60.5) |  |
| Stage II | 84 (24.8) | 7 (18.9) | 8 (21.1) |  |
| Stage III+ | 85 (25.1) | 17 (46.0) | 7 (18.4) |  |
| Pathologic T (%) |  |  |  | 0.0202 |
| T1 | 180 (49.9) | 13 (34.2) | 23 (60.5) |  |
| T2 | 91 (25.2) | 7 (18.4) | 8 (21.1) |  |
| T3 or TX | 90 (24.9) | 18 (47.4) | 7 (18.4) |  |
| Pathologic N (%) |  |  |  | 0.7543 |
| N0 | 246 (68.0) | 31 (81.6) | 33 (86.8) |  |
| NX or N1 | 116 (32.0) | 7 (18.4) | 5 (13.2) |  |
| Pathologic M (%) |  |  |  | 0.0816 |
| M0 | 262 (72.2) | 27 (71.1) | 34 (89.5) |  |
| MX or M1 | 101 (27.8) | 11 (29.0) | 4 (10.5) |  |
| ALB, g/dL | 4.0(3.5, 4.3) | 3.5(3.1, 3.9) | 4.1(3.7, 4.3) | 0.0261 |
| Creatinine, mg/dl | 0.9(0.7, 1.1) | 1.1(0.9, 1.2) | 1.1(0.9, 1.2) | 0.7339 |
| AFP, IU/mL | 15.0(4.0, 264.8) | 27.0(7.0, 779.0) | 7.0(3.0, 261.2) | 0.1141 |
| Platelet count | 211.0(161.0, 299.0) | 197.0 (112.5, 257.8) | 209.0 (170.0, 275.5) | 0.5280 |
| Prothrombin time, seconds | 1.1(1.0, 9.2) | 1.1(0.9, 1.2) | 1.1(1.0, 8.9) | 0.2952 |

* Compared between super and poor survivors.

**Supplementary Table 2. List of top 200 genes with expression level changes in super versus poor survivors**

| Gene | log2FoldChange | pvalue | padj | Regulation |
| --- | --- | --- | --- | --- |
| PADI1 | 5.26 | 2.29E-09 | 4.75E-06 | Up |
| CA9 | -4.22 | 3.98E-10 | 1.41E-06 | Down |
| SLC34A2 | -4.20 | 1.44E-09 | 3.37E-06 | Down |
| MMP10 | -3.88 | 1.73E-12 | 3.23E-08 | Down |
| PNCK | -3.77 | 2.42E-07 | 1.33E-04 | Down |
| CHODL | -3.67 | 2.81E-11 | 1.75E-07 | Down |
| MCOLN3 | -3.55 | 5.90E-10 | 1.58E-06 | Down |
| LOC102724908 | -3.47 | 3.70E-07 | 1.82E-04 | Down |
| CTSV | -3.44 | 7.78E-11 | 3.63E-07 | Down |
| LYPD6B | -3.39 | 1.62E-06 | 4.05E-04 | Down |
| TKTL1 | -3.39 | 3.56E-09 | 6.04E-06 | Down |
| CLEC1B | 3.39 | 8.60E-07 | 2.92E-04 | Up |
| PAGE5 | -3.30 | 2.42E-08 | 3.01E-05 | Down |
| DLK1 | -3.20 | 8.50E-04 | 1.99E-02 | Down |
| SLC1A7 | -3.11 | 4.32E-08 | 4.03E-05 | Down |
| CYP19A1 | -3.10 | 5.40E-06 | 8.34E-04 | Down |
| GNRH2 | 3.04 | 2.02E-07 | 1.18E-04 | Up |
| LAMP5 | -3.04 | 8.35E-08 | 6.78E-05 | Down |
| B3GALT5 | -3.02 | 7.23E-07 | 2.61E-04 | Down |
| SBK3 | -2.91 | 3.09E-09 | 5.76E-06 | Down |
| PEBP4 | 2.87 | 8.53E-06 | 1.10E-03 | Up |
| BARX1 | -2.82 | 3.85E-06 | 6.71E-04 | Down |
| LINC00942 | -2.79 | 1.27E-03 | 2.55E-02 | Down |
| CYP26B1 | -2.76 | 9.28E-09 | 1.33E-05 | Down |
| PLD5 | -2.76 | 2.13E-06 | 4.78E-04 | Down |
| DCHS2 | -2.76 | 1.38E-07 | 1.02E-04 | Down |
| IGHV4-59 | 2.72 | 2.02E-06 | 4.59E-04 | Up |
| IGHV4-28 | 2.72 | 2.33E-05 | 2.08E-03 | Up |
| VEPH1 | -2.72 | 3.94E-08 | 3.87E-05 | Down |
| HOXD10 | -2.72 | 1.81E-06 | 4.34E-04 | Down |
| ENPP6 | -2.71 | 1.29E-07 | 1.00E-04 | Down |
| ADGRG5 | 2.68 | 9.97E-09 | 1.33E-05 | Up |
| SGPP2 | -2.64 | 1.05E-06 | 3.13E-04 | Down |
| IGHV3-74 | 2.63 | 3.54E-06 | 6.42E-04 | Up |
| IGHV1-69 | 2.61 | 2.24E-04 | 8.64E-03 | Up |
| TFF2 | -2.61 | 3.49E-03 | 4.52E-02 | Down |
| CLEC4G | 2.61 | 1.06E-06 | 3.13E-04 | Up |
| RSPO4 | -2.59 | 5.22E-06 | 8.16E-04 | Down |
| ADORA1 | -2.58 | 4.80E-08 | 4.27E-05 | Down |
| GABBR2 | 2.58 | 2.66E-04 | 9.63E-03 | Up |
| CXCL14 | 2.58 | 1.06E-05 | 1.23E-03 | Up |
| ATP10B | -2.55 | 9.60E-05 | 5.18E-03 | Down |
| LCNL1 | -2.55 | 5.55E-04 | 1.56E-02 | Down |
| IGLC3 | 2.53 | 6.40E-06 | 8.98E-04 | Up |
| DLX5 | -2.52 | 5.90E-05 | 3.67E-03 | Down |
| SYN2 | -2.49 | 1.06E-06 | 3.13E-04 | Down |
| IL20RA | -2.49 | 1.64E-03 | 2.91E-02 | Down |
| HMGA2 | -2.49 | 9.10E-06 | 1.13E-03 | Down |
| C3orf52 | -2.49 | 5.03E-07 | 2.27E-04 | Down |
| MROH2B | 2.49 | 4.70E-05 | 3.16E-03 | Up |
| SLC5A1 | 2.47 | 7.19E-05 | 4.26E-03 | Up |
| KRT80 | -2.47 | 5.63E-07 | 2.34E-04 | Down |
| MSC | -2.47 | 5.25E-07 | 2.28E-04 | Down |
| PPFIA4 | -2.46 | 2.79E-08 | 3.25E-05 | Down |
| SPAG17 | -2.44 | 1.23E-06 | 3.41E-04 | Down |
| LINC02982 | -2.43 | 1.08E-05 | 1.24E-03 | Down |
| PRSS22 | -2.43 | 7.65E-04 | 1.86E-02 | Down |
| EPO | -2.42 | 4.23E-04 | 1.31E-02 | Down |
| CLDN18 | -2.41 | 4.84E-05 | 3.21E-03 | Down |
| IGLC2 | 2.40 | 3.41E-06 | 6.30E-04 | Up |
| GALNT17 | 2.39 | 1.26E-04 | 6.14E-03 | Up |
| TNFRSF17 | 2.39 | 7.98E-06 | 1.06E-03 | Up |
| IGHV3-21 | 2.37 | 5.33E-05 | 3.36E-03 | Up |
| UGT8 | -2.36 | 4.98E-04 | 1.47E-02 | Down |
| FHOD3 | -2.36 | 2.43E-06 | 5.17E-04 | Down |
| IGLL5 | 2.36 | 1.24E-05 | 1.37E-03 | Up |
| TWIST2 | -2.35 | 2.42E-05 | 2.13E-03 | Down |
| IGHV3-49 | 2.35 | 1.68E-05 | 1.74E-03 | Up |
| ART5 | -2.35 | 1.01E-05 | 1.21E-03 | Down |
| SPESP1 | -2.35 | 2.04E-05 | 1.98E-03 | Down |
| IGLV3-19 | 2.35 | 4.29E-05 | 2.98E-03 | Up |
| TRIM54 | -2.34 | 4.99E-05 | 3.25E-03 | Down |
| DDN | -2.32 | 5.25E-06 | 8.16E-04 | Down |
| IGHV3-23 | 2.32 | 9.48E-05 | 5.16E-03 | Up |
| SLC7A10 | -2.32 | 1.08E-03 | 2.27E-02 | Down |
| DLGAP2 | 2.30 | 1.95E-04 | 7.95E-03 | Up |
| KHDC1 | -2.27 | 3.51E-06 | 6.42E-04 | Down |
| C5orf58 | -2.27 | 3.32E-05 | 2.58E-03 | Down |
| EREG | -2.26 | 2.41E-04 | 9.07E-03 | Down |
| FRAS1 | -2.26 | 1.85E-05 | 1.85E-03 | Down |
| MUC1 | -2.25 | 1.90E-07 | 1.18E-04 | Down |
| LY6K | -2.25 | 4.38E-06 | 7.37E-04 | Down |
| HTR2B | 2.24 | 9.84E-07 | 3.12E-04 | Up |
| IGHV3-15 | 2.24 | 1.07E-04 | 5.50E-03 | Up |
| CRHBP | 2.24 | 2.23E-07 | 1.26E-04 | Up |
| TFAP2C | -2.23 | 1.80E-03 | 3.05E-02 | Down |
| BPIFB2 | 2.22 | 1.94E-03 | 3.19E-02 | Up |
| IGHV2-70 | 2.21 | 7.28E-04 | 1.81E-02 | Up |
| ITGB8 | -2.21 | 1.34E-04 | 6.29E-03 | Down |
| RCOR2 | -2.20 | 1.32E-05 | 1.43E-03 | Down |
| OVOL2 | -2.19 | 3.98E-04 | 1.25E-02 | Down |
| CTSG | 2.19 | 4.84E-05 | 3.21E-03 | Up |
| KCNJ16 | -2.17 | 6.90E-04 | 1.76E-02 | Down |
| IGHV1-2 | 2.17 | 9.21E-04 | 2.08E-02 | Up |
| C1QL4 | -2.16 | 1.63E-04 | 7.15E-03 | Down |
| ZNF860 | -2.16 | 1.88E-06 | 4.39E-04 | Down |
| SCIN | -2.15 | 8.30E-09 | 1.29E-05 | Down |
| FCN2 | 2.15 | 2.49E-04 | 9.25E-03 | Up |
| LINC02331 | -2.15 | 2.95E-04 | 1.04E-02 | Down |
| LOC644135 | -2.14 | 6.48E-06 | 9.03E-04 | Down |
| FCN3 | 2.13 | 5.11E-07 | 2.27E-04 | Up |
| MROH2A | 2.12 | 2.39E-04 | 9.06E-03 | Up |
| ADAM23 | -2.11 | 4.44E-05 | 3.06E-03 | Down |
| FSIP1 | -2.10 | 2.98E-08 | 3.27E-05 | Down |
| NOC2LP1 | -2.09 | 2.67E-06 | 5.60E-04 | Down |
| SPOCK1 | -2.08 | 1.80E-05 | 1.83E-03 | Down |
| MMP7 | -2.08 | 6.89E-04 | 1.76E-02 | Down |
| IGFALS | 2.08 | 5.92E-06 | 8.64E-04 | Up |
| S100A1 | 2.07 | 7.03E-07 | 2.61E-04 | Up |
| CLEC17A | 2.06 | 3.91E-05 | 2.82E-03 | Up |
| IGLV1-44 | 2.05 | 3.44E-04 | 1.13E-02 | Up |
| IGKV1-17 | 2.04 | 1.60E-03 | 2.87E-02 | Up |
| NPTXR | -2.04 | 3.82E-06 | 6.71E-04 | Down |
| CCDC74B | -2.04 | 1.89E-05 | 1.88E-03 | Down |
| COL22A1 | 2.04 | 1.45E-03 | 2.73E-02 | Up |
| ASS1P11 | 2.04 | 9.68E-06 | 1.18E-03 | Up |
| CXCL1 | -2.03 | 1.62E-04 | 7.15E-03 | Down |
| DIO2 | 2.03 | 1.34E-04 | 6.29E-03 | Up |
| PZP | 2.02 | 2.74E-05 | 2.26E-03 | Up |
| NCAPD2P1 | -2.02 | 9.28E-04 | 2.09E-02 | Down |
| CALCR | -2.02 | 9.91E-05 | 5.27E-03 | Down |
| FRRS1L | 2.00 | 1.46E-03 | 2.73E-02 | Up |
| PKIA | -2.00 | 4.50E-05 | 3.08E-03 | Down |
| TMEM132B | -2.00 | 1.48E-05 | 1.56E-03 | Down |
| FER1L5 | -1.99 | 9.59E-07 | 3.09E-04 | Down |
| IGLV1-40 | 1.99 | 2.45E-04 | 9.11E-03 | Up |
| IGHG1 | 1.99 | 3.02E-04 | 1.05E-02 | Up |
| AOC1 | -1.98 | 2.17E-04 | 8.46E-03 | Down |
| LOC101927793 | 1.98 | 3.25E-04 | 1.10E-02 | Up |
| RIC3 | 1.98 | 1.49E-04 | 6.67E-03 | Up |
| FLNC | -1.98 | 1.83E-04 | 7.64E-03 | Down |
| NSG1 | 1.97 | 7.49E-07 | 2.64E-04 | Up |
| COLEC10 | 1.97 | 5.78E-06 | 8.64E-04 | Up |
| EFNA5 | -1.96 | 1.19E-04 | 5.93E-03 | Down |
| LINC01093 | 1.96 | 1.43E-04 | 6.53E-03 | Up |
| C5orf46 | -1.96 | 7.78E-05 | 4.47E-03 | Down |
| TREM1 | -1.95 | 1.63E-06 | 4.05E-04 | Down |
| DPT | 1.95 | 3.29E-04 | 1.10E-02 | Up |
| IGHV1-69D | 1.95 | 1.90E-03 | 3.16E-02 | Up |
| TERT | 1.94 | 2.86E-04 | 1.01E-02 | Up |
| KCNH2 | -1.93 | 6.20E-04 | 1.67E-02 | Down |
| NPTX1 | -1.93 | 1.66E-04 | 7.18E-03 | Down |
| CDH17 | -1.92 | 8.06E-04 | 1.92E-02 | Down |
| SH2D5 | -1.92 | 1.37E-06 | 3.50E-04 | Down |
| ZNF804A | -1.92 | 7.94E-05 | 4.55E-03 | Down |
| GOLGA7B | -1.92 | 8.83E-05 | 4.92E-03 | Down |
| TMPRSS4 | -1.91 | 3.87E-04 | 1.22E-02 | Down |
| RAET1K | -1.91 | 4.91E-06 | 7.90E-04 | Down |
| ZNF215 | -1.90 | 1.24E-06 | 3.41E-04 | Down |
| SPP1 | -1.90 | 2.59E-04 | 9.43E-03 | Down |
| IGKV1-6 | 1.90 | 1.10E-03 | 2.30E-02 | Up |
| DNASE1L3 | 1.90 | 4.52E-10 | 1.41E-06 | Up |
| DIO3OS | 1.89 | 3.25E-04 | 1.10E-02 | Up |
| CERS1 | -1.89 | 5.92E-04 | 1.62E-02 | Down |
| KLC3 | -1.89 | 8.66E-04 | 2.02E-02 | Down |
| ELOVL4 | -1.89 | 3.57E-05 | 2.69E-03 | Down |
| CEL | 1.88 | 1.10E-04 | 5.60E-03 | Up |
| CCL25 | 1.88 | 6.33E-04 | 1.68E-02 | Up |
| ELFN1 | 1.88 | 6.61E-06 | 9.08E-04 | Up |
| MEGF10 | 1.88 | 1.17E-04 | 5.88E-03 | Up |
| CDCP1 | -1.88 | 1.61E-05 | 1.68E-03 | Down |
| GUCY2C | -1.88 | 2.10E-03 | 3.34E-02 | Down |
| GAPLINC | 1.88 | 3.74E-04 | 1.20E-02 | Up |
| AJAP1 | -1.87 | 1.58E-04 | 7.03E-03 | Down |
| ANXA10 | 1.87 | 5.67E-06 | 8.60E-04 | Up |
| LINC02761 | 1.86 | 8.38E-06 | 1.09E-03 | Up |
| SCT | 1.86 | 1.35E-05 | 1.45E-03 | Up |
| MYEF2 | -1.86 | 1.62E-05 | 1.68E-03 | Down |
| CDH16 | -1.85 | 6.26E-04 | 1.68E-02 | Down |
| LOC100419170 | -1.85 | 7.25E-04 | 1.80E-02 | Down |
| LINC00402 | 1.85 | 6.31E-04 | 1.68E-02 | Up |
| MBOAT4 | -1.85 | 1.22E-05 | 1.35E-03 | Down |
| CREG2 | -1.84 | 1.97E-07 | 1.18E-04 | Down |
| TMEM100 | 1.84 | 3.89E-05 | 2.82E-03 | Up |
| FAM225A | -1.84 | 1.27E-05 | 1.39E-03 | Down |
| CADPS | -1.83 | 3.80E-03 | 4.74E-02 | Down |
| MATN3 | -1.83 | 1.35E-04 | 6.30E-03 | Down |
| ZNF488 | -1.83 | 2.55E-04 | 9.37E-03 | Down |
| LVRN | -1.83 | 2.47E-05 | 2.13E-03 | Down |
| EPHB6 | -1.83 | 2.76E-05 | 2.26E-03 | Down |
| LINC02490 | 1.82 | 6.67E-05 | 4.05E-03 | Up |
| MYCN | -1.82 | 3.54E-04 | 1.15E-02 | Down |
| TMEM132A | -1.82 | 1.30E-06 | 3.42E-04 | Down |
| ACP3 | -1.82 | 3.90E-04 | 1.23E-02 | Down |
| CELF4 | -1.82 | 1.74E-05 | 1.78E-03 | Down |
| FCRL5 | 1.82 | 2.89E-04 | 1.02E-02 | Up |
| EGLN3 | -1.82 | 4.56E-06 | 7.60E-04 | Down |
| PFKP | -1.81 | 3.05E-07 | 1.54E-04 | Down |
| FAM99A | 1.81 | 1.96E-03 | 3.19E-02 | Up |
| NCAM1 | -1.81 | 1.42E-04 | 6.53E-03 | Down |
| RPS20P6 | 1.80 | 1.22E-04 | 6.04E-03 | Up |
| LOC102723834 | -1.80 | 3.20E-05 | 2.52E-03 | Down |
| PTK7 | -1.80 | 2.09E-05 | 1.99E-03 | Down |
| IGHV3-43 | 1.80 | 1.96E-03 | 3.20E-02 | Up |
| TTC36 | 1.79 | 9.69E-04 | 2.15E-02 | Up |
| SLC4A10 | 1.79 | 2.40E-05 | 2.12E-03 | Up |
| CNTFR | 1.78 | 2.41E-03 | 3.61E-02 | Up |
| PNOC | 1.78 | 6.95E-04 | 1.76E-02 | Up |
| REN | 1.78 | 4.22E-04 | 1.31E-02 | Up |
| MMP12 | -1.78 | 1.30E-03 | 2.58E-02 | Down |

**Supplementary Table 3. Reported function of the 19 selected signature genes involved in immune response, metabolism, and cell cycle processes**

| Gene | Function |
| --- | --- |
| Genes identified through immune response pathways | |
| CCNA2 | CCNA2 (Cyclin A2) is a Protein Coding gene, which belongs to the highly conserved cyclin family, whose members function as regulators of the cell cycle. This protein binds and activates cyclin-dependent kinase 2 and thus promotes transition through G1/S and G2/M. |
| CD8A | The CD8 antigen is a cell surface glycoprotein found on most cytotoxic T lymphocytes that mediates efficient cell-cell interactions within the immune system. The CD8 antigen acts as a coreceptor with the T-cell receptor on the T lymphocyte to recognize antigens displayed by an antigen presenting cell in the context of class I MHC molecules. |
| CNR2 | Cannabinoid Receptor 2, also known as the peripheral cannabinoid receptor, is a member of the cannabinoid receptor group of G-protein-coupled receptors that also includes CB1 and GPR55. Cannabinoid receptors 2 are located primarily in immune cells both within and outside the CNS. |
| CTSG | Cathepsin G, a member of the peptidase S1 protein family, is found in azurophil granules of neutrophilic polymorphonuclear leukocytes. The encoded protease has a specificity similar to that of chymotrypsin C, and may participate in the killing and digestion of engulfed pathogens, and in connective tissue remodeling at sites of inflammation. In addition, the encoded protein is antimicrobial, with bacteriocidal activity against S. aureus and N. gonorrhoeae. |
| HSPA6 | Heat Shock Protein Family A (Hsp70) Member 6, involved in cellular response to heat and protein refolding. Located in centriole and cytosol. Colocalizes with COP9 signalosome |
| ITGA2B | Integrin Subunit Alpha 2b. This gene encodes a member of the integrin alpha chain family of proteins. The encoded preproprotein is proteolytically processed to generate light and heavy chains that associate through disulfide linkages to form a subunit of the alpha-IIb/beta-3 integrin cell adhesion receptor. This receptor plays a crucial role in the blood coagulation system, by mediating platelet aggregation. Mutations in this gene are associated with platelet-type bleeding disorders, which are characterized by a failure of platelet aggregation, including Glanzmann thrombasthenia. |
| RTKN2 | Involved in negative regulation of intrinsic apoptotic signaling pathway; positive regulation of NF-kappaB transcription factor activity; and positive regulation of NIK/NF-kappaB signaling. Located in cytoplasm and nucleus. May play an important role in lymphopoiesis. |
| S100A9 | S100A9 is a calcium- and zinc-binding protein which plays a prominent role in the regulation of inflammatory processes and immune response. It can induce neutrophil chemotaxis, adhesion, can increase the bactericidal activity of neutrophils by promoting phagocytosis via activation of SYK, PI3K/AKT, and ERK1/2 and can induce degranulation of neutrophils by a MAPK-dependent mechanism. |
| SPP1 | This protein is a cytokine that upregulates expression of interferon-gamma and interleukin-12. This protein might also reduce production of interleukin-10 and is essential in the pathway that leads to type I immunity. |
| Genes identified through metabolism pathways | |
| EGLN3 | Egl-9 Family Hypoxia Inducible Factor 3. Involved in several processes, including activation of cysteine-type endopeptidase activity involved in apoptotic process; peptidyl-proline hydroxylation to 4-hydroxy-L-proline; and response to hypoxia. Located in cytosol and nucleus. Implicated in renal cell carcinoma. |
| ENO2 | Enolase 2. Diseases associated with ENO2 include Glycogen Storage Disease Due To Muscle Beta-Enolase Deficiency and Granular Cell Tumor. Among its related pathways are glycolysis (BioCyc) and Gluconeogenesis. Gene Ontology (GO) annotations related to this gene include magnesium ion binding and phosphopyruvate hydratase activity. |
| KIF20A | Kinesin Family Member 20A. Among its related pathways are Golgi-to-ER retrograde transport and Cell Cycle, Mitotic. Gene Ontology (GO) annotations related to this gene include protein kinase binding and ATP hydrolysis activity. |
| PKM | Pyruvate Kinase M1/2. This gene encodes a protein involved in glycolysis. The encoded protein is a pyruvate kinase that catalyzes the transfer of a phosphoryl group from phosphoenolpyruvate to ADP, generating ATP and pyruvate. This protein has been shown to interact with thyroid hormone and may mediate cellular metabolic effects induced by thyroid hormones. |
| RDH16 | Enables NAD-retinol dehydrogenase activity; androstan-3-alpha,17-beta-diol dehydrogenase activity; and androsterone dehydrogenase activity. Involved in steroid metabolic process. Located in intracellular membrane-bounded organelle |
| SLC16A3 | Solute Carrier Family 16 Member 3. Gene Ontology (GO) annotations related to this gene include RNA binding and monocarboxylic acid transmembrane transporter activity. Proton-dependent transporter of monocarboxylates such as L-lactate and pyruvate. Plays a predominant role in L-lactate efflux from highly glycolytic cells. |
| Genes identified through cell cycle pathways | |
| CD44 | CD44 Molecule. The protein encoded by this gene is a cell-surface glycoprotein involved in cell-cell interactions, cell adhesion and migration. It is a receptor for hyaluronic acid (HA) and can also interact with other ligands, such as osteopontin, collagens, and matrix metalloproteinases (MMPs). This protein participates in a wide variety of cellular functions including lymphocyte activation, recirculation and homing, hematopoiesis, and tumor metastasis. |
| CXCL1 | C-X-C Motif Chemokine Ligand 1. This antimicrobial gene encodes a member of the CXC subfamily of chemokines. The encoded protein is a secreted growth factor that signals through the G-protein coupled receptor, CXC receptor 2. This protein plays a role in inflammation and as a chemoattractant for neutrophils. Aberrant expression of this protein is associated with the growth and progression of certain tumors. |
| SPOCK1 | This gene encodes the protein core of a seminal plasma proteoglycan containing chondroitin- and heparan-sulfate chains. The protein's function is unknown, although similarity to thyropin-type cysteine protease-inhibitors suggests its function may be related to protease inhibition. May play a role in cell-cell and cell-matrix interactions. May contribute to various neuronal mechanisms in the central nervous system. |
| TTK | TTK Protein Kinase. This gene encodes a dual specificity protein kinase with the ability to phosphorylate tyrosine, serine and threonine. Associated with cell proliferation, this protein is essential for chromosome alignment at the centromere during mitosis and is required for centrosome duplication. It has been found to be a critical mitotic checkpoint protein for accurate segregation of chromosomes during mitosis. Tumorigenesis may occur when this protein fails to degrade and produces excess centrosomes resulting in aberrant mitotic spindles. |

Note: The reported gene functions were retrieved from <https://www.genecards.org/>.

**Supplementary Table 4. Prognostic gene signatures for HCC survival with external validation from literature review**

| Ref. | Gene signature | Methods of identifying DEGs | Sample size in discovery cohort | Data source | | | AUROC in discovery cohort | AUROC in validation cohort | |
| --- | --- | --- | --- | --- | --- | --- | --- | --- | --- |
| 1 | 5 metastasis-related gene signature | HCC vs normal tissues | 233 | TCGA (D)  ICGC (V) | | | 1-, 2-, 3-year  0.786,0.786 and 0.777 | 1-, 2-, 3-year  0.703, 709, 713 | |
| 2 | 10 Ferroptosis-related gene signature | HCC vs normal tissues | 365 | TCGA (D)  ICGC (V) | | | 1-, 3-, 5-year  0.800, 0.690, 0.668 | 1-, 3-, 5-year  0.680, 0.690 0.718 | |
| 3 | 6 gene signature | HCC vs normal tissues | 343 | TCGA (D)  GSE14520 (V) | | | 1-, 3-, 5-year  0.773, 0.702, 0.673 | 1-, 3-, 5-year  0.678, 0.643, and 0.633 | |
| 4 | 10 peroxisome-related gene signature | LASSO-COX | 351 | TCGA (D)  ICGC (V) | | | 1-, 3-, 5-year  0.715, 0.704, 0.691 | 1-, 3-, 5-year  0.759, 0.783, 0.724 | |
| 5 | 8 cuproptosis-related gene signature | Two clusters from K-means analysis | 363 | TCGA (D)  GSE14520, Pan-Cancer Analysis of Whole Genomes (V) | | | 1-, 3-, 5-year  0.756, 0.706, and 0.722 | 1-, 3-, 5-year  0.744， 0.715， 0.757 | |
| 6 | 14 redox-immune related gene signature | LASSO-COX | 116 | TCGA (D)  ICGC (V)  GSE14520 (V) | | | 1-, 3-, 5-year  0.802, 793, 0.755 | 1-, 3-, 5-year  ICGC 0.750, 0.668, 0.708;  GSE 0.678, 0.727, 0.708 | |
| 7 | 3 metabolic rate-limiting enzymes related gene signature | Mouse liver cancer tissues vs paired normal tissues;  Cox | Train: 116  Test: 119 | TCGA (D)  HCC patients tissue microarray by IHC (V) | | | 1-, 3-, 5-year  Train: 0.745, 0.71, 0732  Test:0.84, 0.85, 0754 | 1-, 3-, 5-year  0.767, 0.744, 0.803 | |
| 8 | 10 metabolic gene signature | Cox | 342 | TCGA (D)  GSE14520 (V)  ICGC (V)  Clinical cohort (V) | | | 3-, 5-year  0.750, 0.722 (gene signature plus TNM stage) | 3-, 5-year  GSE 0.727, 0.768  ICGC 0.743, 0.572  Clinical cohort 0.735,0.701 (gene signature plus TNM stage) | |
| 9 | 7 senescence-associated gene signature | COX | 209 | | GSE14520 (D)  TCGA (V) | Not reported | | | 1-,3-,5-year  0.708, 0.69, 0.678 |
| 10 | 4 cuproptosis-related gene signature | LASSO-COX | 365 | | TCGA (D)  ICGC (V) | 1-, 2-, 3-year  0.734, 0.659, 0.646 | | | 1-, 2-, 3-year  0.588, 0.651, 0.677 |
| 11 | 4 macrophages-related gene signature | HCC vs normal tissues | 351 | | TCGA (D)  GSE14520 (V) | 1-, 3-, 5-year  0.798, 0.748, 0.721 | | | 1-, 3-, 5-year  0.654, 0.608, 0.625 |
| 12 | 4 gene signature based on nonsense-mediated RNA decay | LASSO-COX | Not reported | | TCGA (D)  GSE54236, GSE116174, GSE76427 (V) | 2-, 3-, 4-year  0.681, 0.687, 0.702 | | | 2-, 3-, 4-year  0.577, 0.637, 0.638 |
| 13 | 10-immune related gene signature | LASSO-COX | 364 | | TCGA (D)  GSE14520 (V)  ICGC (V) | 1-, 2-, 3-year  0.739, 0.731, 0.695 | | | 1-, 2-, 3-year  GSE 0.667, 0.702, 0.675  ICGC 0.747, 0.752, 0.764 |
| 14 | 5 EMT‑related gene signature | COX | 319 | | TCGA (D)  ICGC (V) | 1-, 2-, 3-year  0.803, 0.721, 0.7 | | | 1-, 2-, 3-year  0.739, 0.74, 0.754 |

D: discovery cohort; V: validation cohort

**Supplementary Figure 1. Misclassifications on super and poor survivors by different gene signatures in matched HCC patients.**

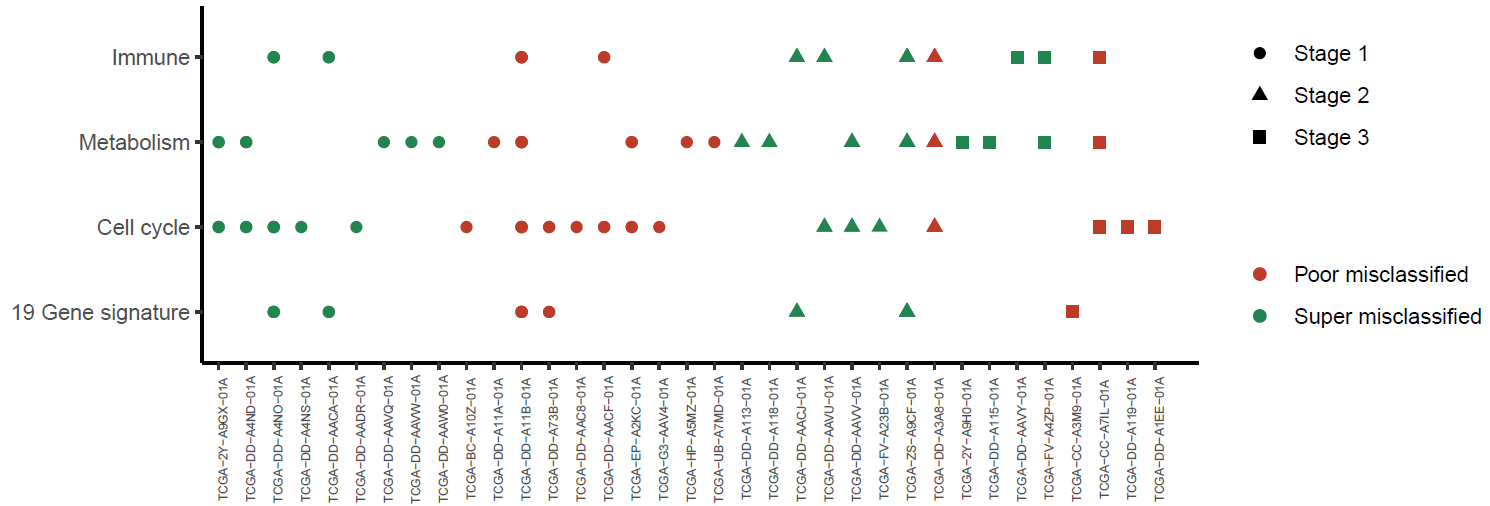

**Supplementary Figure 2.** **Clinical performance of the combined gene signatures together with clinical factors.**

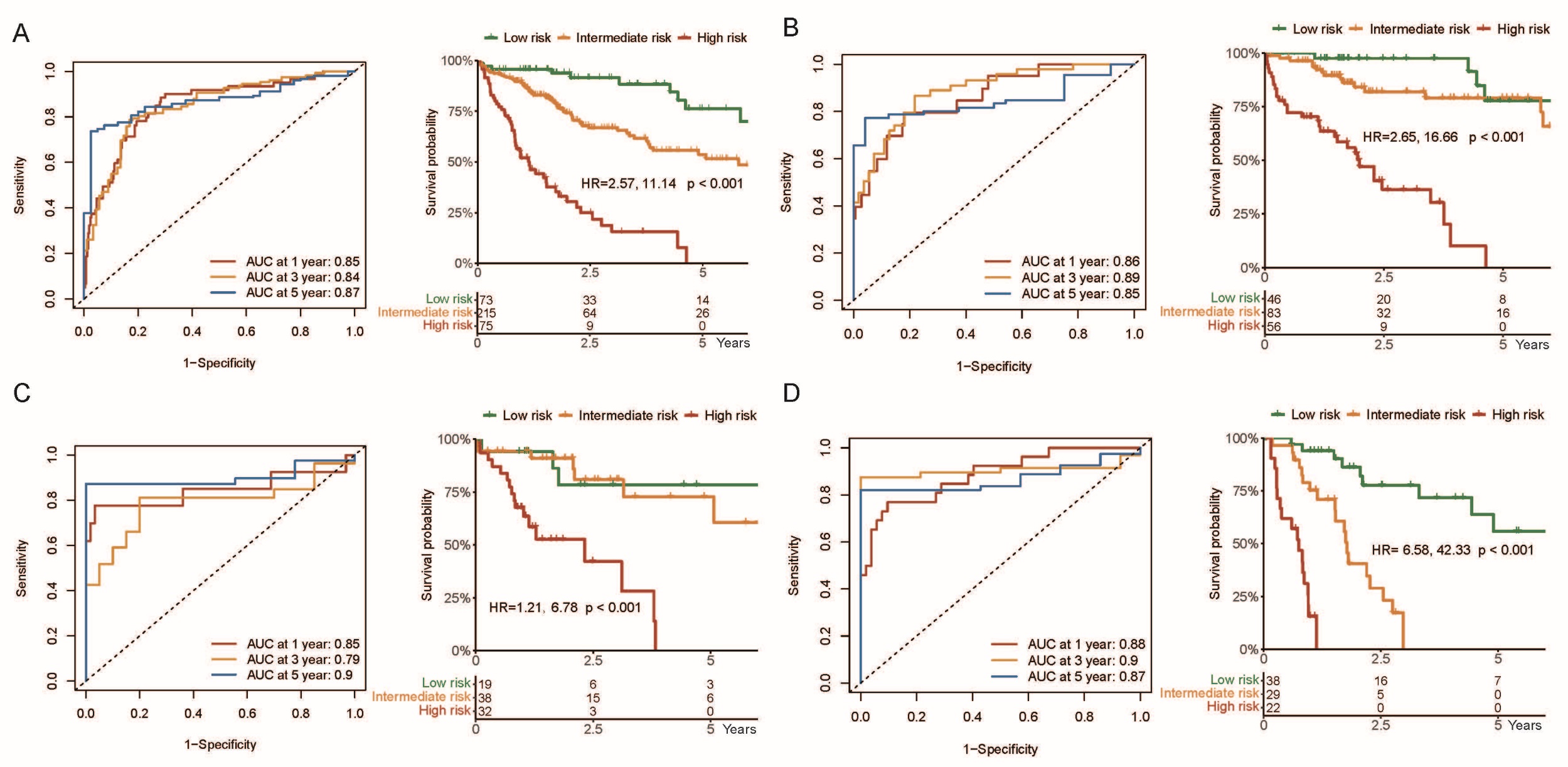

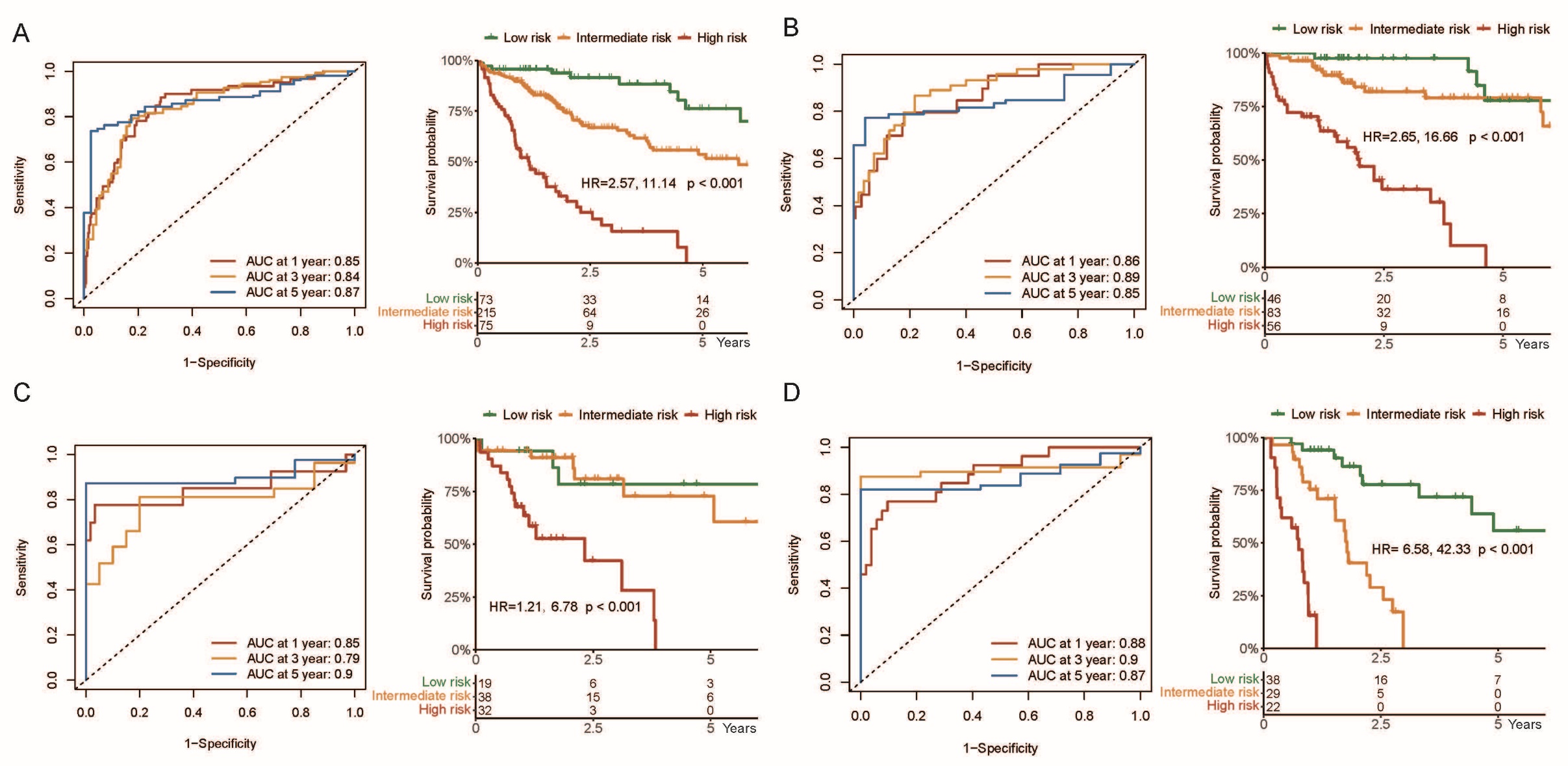

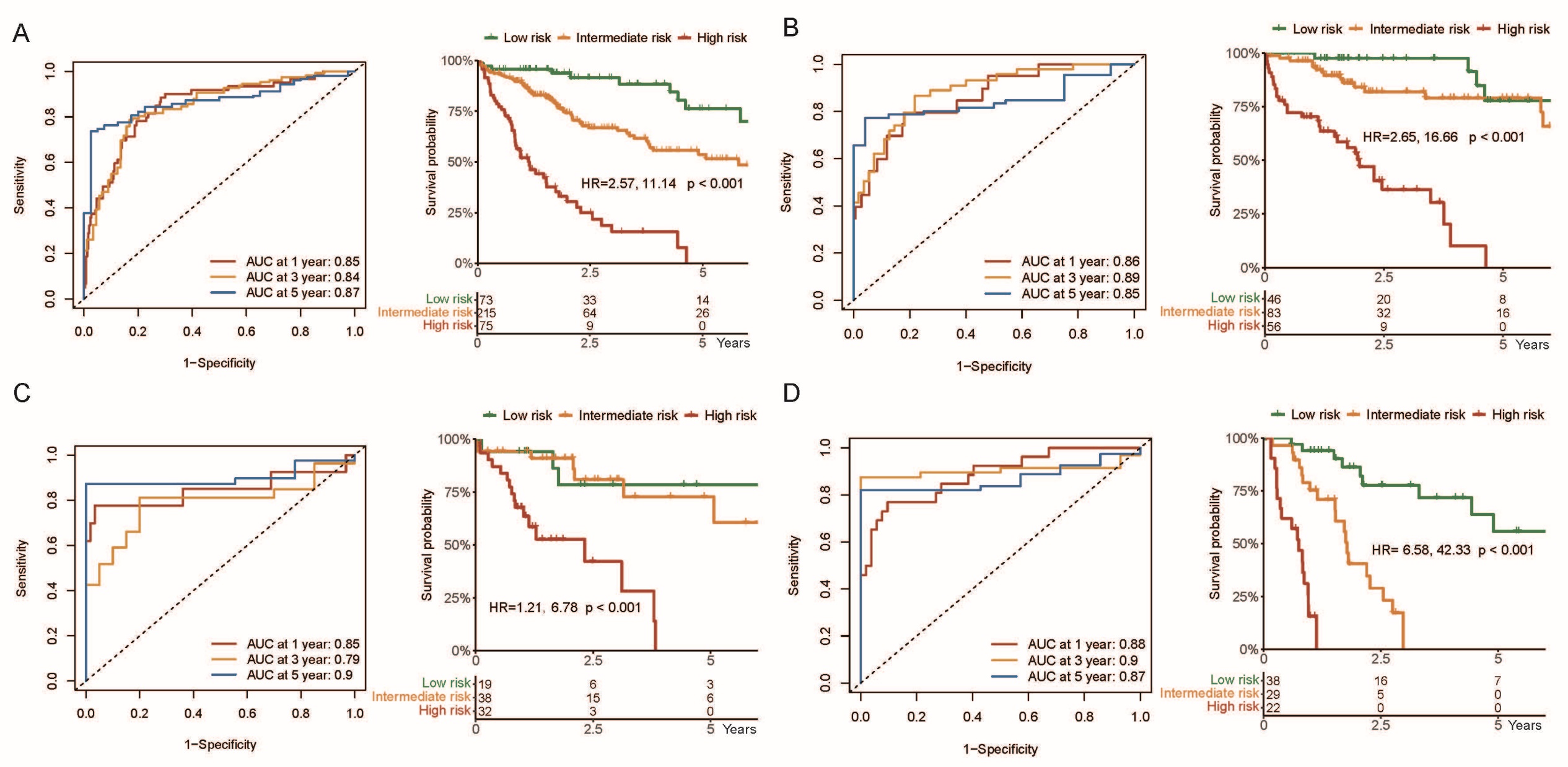

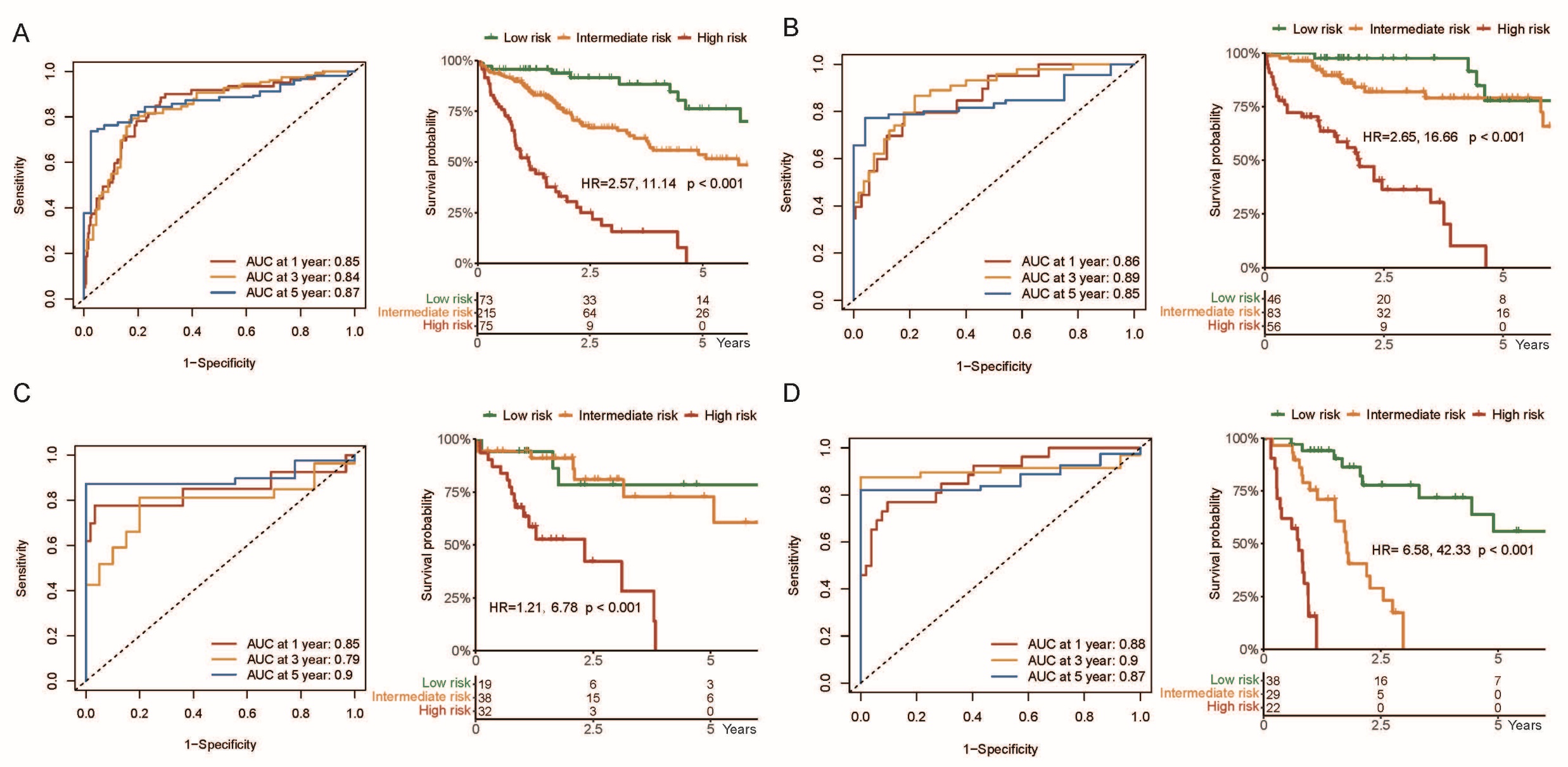

AUROC and Kaplan-Meier survival curves of combined gene sets and clinical predictors in predicting the overall survival in total HCC patients from TCGA (A), and HCC patients from TCGA in stage Ⅰ (B), Ⅱ (C) and Ⅲ (D). Cutoffs in total HCC patients were determined by 20% percentile (cutoff=-0.8) and 80% (cutoff=1) percentile.
